# GenomicSuperSignature: interpretation of RNA-seq experiments through robust, efficient comparison to public databases

**DOI:** 10.1101/2021.05.26.445900

**Authors:** Sehyun Oh, Ludwig Geistlinger, Marcel Ramos, Daniel Blankenberg, Marius van den Beek, Jaclyn N. Taroni, Vincent Carey, Casey Greene, Levi Waldron, Sean Davis

**Author notes:** equal contribution (Levi Waldron, Sean Davis).

## Abstract

Millions of transcriptomic profiles have been deposited in public archives, yet remain underused for the interpretation of new experiments. We present a novel method for interpreting new transcriptomic datasets through near-instantaneous comparison to public archives without high-performance computing requirements. We performed Principal Component Analysis on 536 studies comprising 44,890 RNA sequencing profiles. Sufficiently similar loading vectors were aggregated to form *R*eplicable *A*xes of *V*ariation (RAV). RAVs were annotated with metadata of originating studies and samples and by gene set enrichment analysis. Functionality to associate new datasets with RAVs, extract interpretable annotations, and provide intuitive visualization are implemented as the GenomicSuperSignature R/Bioconductor package. We demonstrated the efficient and coherent database searching, robustness to batch effects and heterogeneous training data, and transfer learning capacity of our method using TCGA and rare diseases datasets. GenomicSuperSignature will aid analyzing new gene expression data in the context of existing databases using minimal computing resources.

**PURPOSE:** Millions of transcriptomic profiles have been deposited in public archives, yet remain underused for the interpretation of new experiments. Existing methods for leveraging these public resources have focused on the reanalysis of existing data or analysis of new datasets independently. We present a novel approach to interpreting new transcriptomic datasets by near-instantaneous comparison to public archives without high-performance computing requirements.

**METHODS:** To identify replicable and interpretable axes of variation in any given gene expression dataset, we performed Principal Component Analysis (PCA) on 536 studies comprising 44,890 RNA sequencing profiles. Sufficiently similar loading vectors, when compared across studies, were aggregated to form *R*eplicable *A*xes of *V*ariation (RAV). RAVs were annotated with metadata of originating studies and samples and by gene set enrichment analysis. Functionality to associate new datasets with RAVs, extract interpretable annotations, and provide intuitive visualization are implemented as the GenomicSuperSignature R/Bioconductor package.

**RESULTS:** RAVs are robust to batch effects and the presence of low-quality or irrelevant studies, and identify signals that can be lost by merging samples across the training datasets. The GenomicSuperSignature package allows instantaneous matching of PCA axes in new datasets to pre-computed RAVs, cutting down the analysis time from days to the order of seconds on an ordinary laptop. We demonstrate that RAVs associated with a phenotype can provide insight into weak or indirectly measured biological attributes in a new study by leveraging accumulated data from published datasets. Benchmarking against complementary previous works demonstrates that the RAV index 1) identifies colorectal carcinoma transcriptome subtypes that are similar to but more correlated with clinicopathological characteristics than previous disease-specific efforts and 2) can estimate neutrophil counts through transfer learning on new data comparably to the previous efforts despite major differences in training datasets and model building processes with the additional benefits of flexibility and scalability of the model application.

**CONCLUSION:** GenomicSuperSignature establishes an information resource and software tools to interrogate it. Prior knowledge databases are coherently linked, enabling researchers to analyze new gene expression data in the context of existing databases using minimal computing resources. The robustness of GenomicSuperSignature suggests that we can expand this approach beyond human gene expression profiles, such as single-cell RNA-seq, microbiome abundance, and different species’ transcriptomics datasets.

## Introduction

Vast quantities of transcriptomic data have been deposited in public archives, yet remain underused for the interpretation of new experiments. Here, we present a novel toolkit and approach for interpreting new transcriptomic datasets through near-instantaneous comparison to public transcriptomic datasets using the compute resources available on a standard laptop.

Dimensionality reduction has been broadly adopted to transform transcriptomes onto a smaller number of latent variables representing co-expressed transcripts. Gene co-expression can result from shared function or regulation^1^, association with tissue composition or cell type^2^, and technical batch effects^3^. In the confluence of these factors, dimensionality reduction can assist interpretability and reduce the burden of multiple hypothesis testing, but can also lead to incomplete or misleading interpretation. Valid interpretation would be improved by comparison of latent variables in new datasets to those also present in public transcriptome databases.

Many dimensionality reduction approaches, differing in the optimization and constraint criteria, are available^4^ and there have been multiple attempts to detect biological and technical signals through these lower dimensional, latent variable representations. Classic methods such as Principal Component Analysis (PCA) and non-negative matrix factorization (NMF) remain widely used in their original form and as bases for newer methods. For example, scCoGAPS is an NMF method optimized for large, sparse single cell RNA sequencing datasets^5^. scCoGAPS recovers features in a source dataset, then projects a new dataset onto this learned latent space through projectR^5,6^. This approach requires users to train their own model and mostly focuses on single cell RNA sequencing datasets with similar biology. PLIER aims to improve on the interpretability of PCA by identifying latent variables that map to a single gene set or a group of highly related gene sets with positive correlations^7^. MultiPLIER applies the PLIER approach to transfer learned patterns from a large public dataset to rare diseases^8^. Other tools focus on recovering consistent signals from multiple datasets across distinct platforms^9,10^, increasing interpretability^11^, simple database search^12^, or transfer learning between datasets of a specific type^13,14^. However, none of these tools enable routine exploratory analysis of new studies through comparison to large public transcriptome databases. Also, these tools do not provide a reference catalog for transfer learning from large public databases, or in the case of MultiPLIER, require substantial computing resources and bioinformatics expertise.

Here, we describe GenomicSuperSignature, a toolkit for interpreting new RNA-seq datasets by dimensionality reduction in the context of a large-scale database of previously published and annotated results. As an exploratory data analysis tool, GenomicSuperSignature matches PCA axes in a new dataset to an annotated index of ***r***eplicable ***a***xes of ***v***ariation (RAV) that are represented in previously published independent datasets. GenomicSuperSignature also can be used as a tool for *transfer learning^15^*, utilizing RAVs as well-defined and replicable latent variables defined by multiple previous studies in place of *de novo* latent variables. The interpretability of RAVs is enhanced through annotations by *ME*dical *S*ubject *H*eadings (MeSH) and *G*ene *S*et *E*nrichment *A*nalysis (GSEA). Through the use of pre-built, pre-annotated, dimension-reduced RAVs, GenomicSuperSignature leverages knowledge from tens of thousands of samples and from PubMed and MSigDB^16^, to the dataset at hand within seconds on an ordinary laptop. We demonstrate these functionalities in colorectal carcinoma, breast invasive carcinoma, systemic lupus erythematosus, and rare inflammatory disease. GenomicSuperSignature is implemented as an R/Bioconductor package for straightforward incorporation into popular RNA-seq analysis pipelines.

## Results

The current RAVmodel is trained on 536 studies containing 44,890 human RNA sequencing profiles. This RAVmodel is associated with 18,798 (4,430 unique) MeSH terms and 70,687 (1,784 unique) MSigDB curated (C2) gene sets. This integration of data resources (Fig.1B) is accompanied by tools in the GenomicSuperSignature R/Bioconductor package for interpretation of new datasets (Fig. 1A and Supplementary Fig. 1C). We demonstrate this novel application of public data in three examples. First, using TCGA datasets, we showed that new data can be rapidly associated with related studies, gene sets, and MeSH terms. Second, we showed that the RAVmodel trained from diverse RNA-seq experiments identified colon cancer transcriptome subtypes more closely associated to clinicopathological variables than the subtypes previously identified by meta-analysis of a focused colorectal carcinoma (CRC) microarray compendium. Lastly, we showed that neutrophil counts of two independent datasets can be interpreted and inferred through a single RAV, providing a novel quantitative measure of neutrophil count from transcriptome data. These examples, along with sensitivity analyses and simulations, demonstrate that the RAVmodel and GenomicSuperSignature are a robust, general-purpose method for the interpretation of transcriptome data.

**Fig 1.**
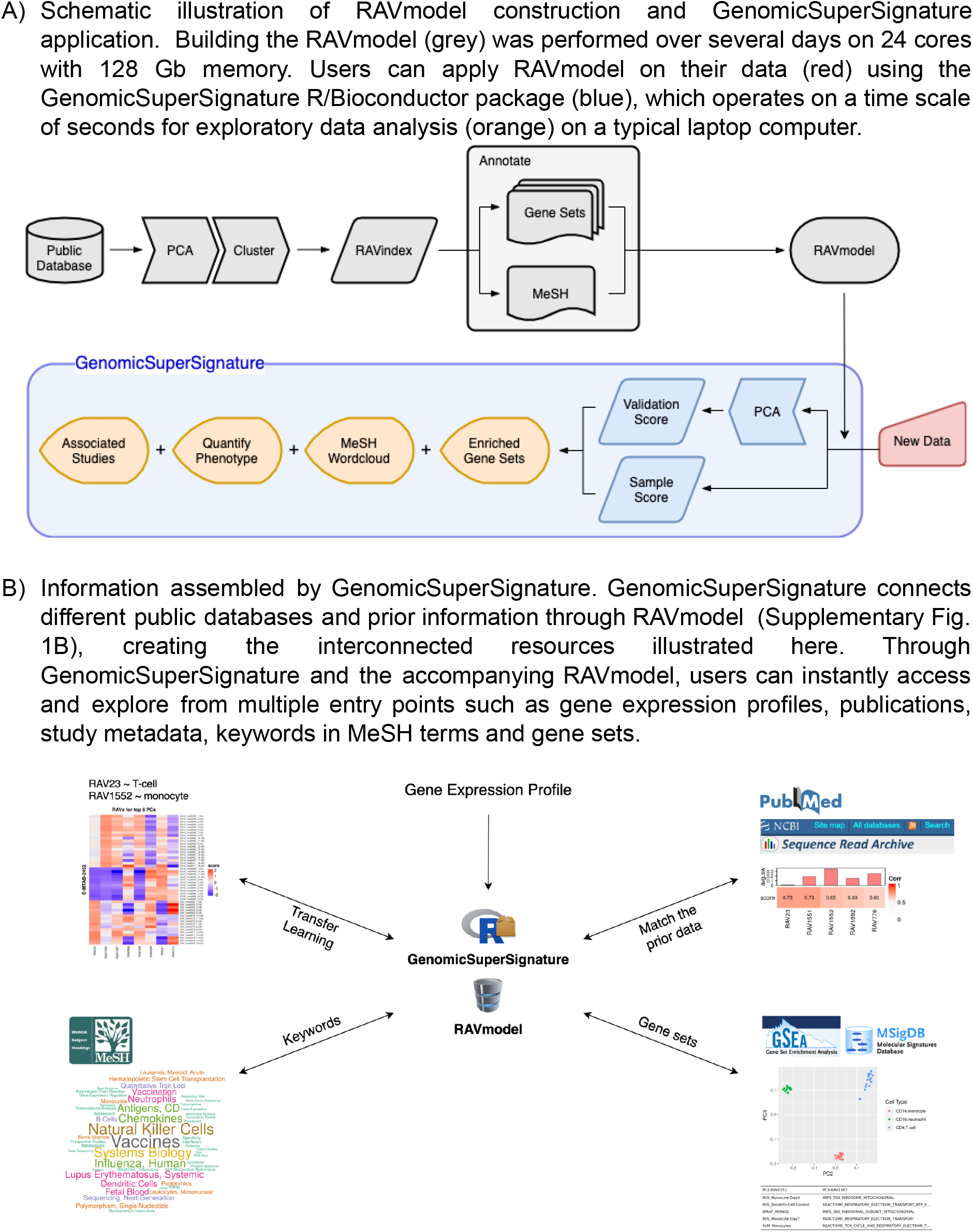
Overview of GenomicSuperSignature. A) Schematic illustration of RAVmodel construction and GenomicSuperSignature application. Building the RAVmodel (grey) was performed over several days on 24 cores with 128 Gb memory. Users can apply RAVmodel on their data (red) using the GenomicSuperSignature R/Bioconductor package (blue), which operates on a time scale of seconds for exploratory data analysis (orange) on a typical laptop computer. B) Information assembled by GenomicSuperSignature. GenomicSuperSignature connects different public databases and prior information through RAVmodel (Supplementary Fig. 1B), creating the interconnected resources illustrated here. Through GenomicSuperSignature and the accompanying RAVmodel, users can instantly access and explore from multiple entry points such as gene expression profiles, publications, study metadata, keywords in MeSH terms and gene sets.

### Sensitivity Analysis and Simulation

Model training methods were optimized for robustness, simplicity, computational cost, and validity (Supplementary Methods). Briefly, the RAVmodel was trained on the RNA-seq Sample Compendia of refine.bio^17^. We analyzed TPM count data using Principal Components Analysis (PCA) following log-transformation, then identified clusters of similar principal components from independent datasets using hierarchical clustering on Spearman distance and ward.D agglomeration. This approach was compared to alternatives based on 1) ability to group synthetic true-positive principal components, 2) separation of synthetic true-negative principal components added to training data, 3) magnitude of changes in the results compared to the simplest method, and 4) maintenance of RAVs identified from a focused training dataset when adding unrelated datasets. Alternative approaches considered but not selected for training the final model included NMF, Independent Component Analysis (ICA), PLIER^18^ and MultiPLIER^8^, Variance-Stabilizing Transformation (VST)^19^ in place of log transformation, combining training datasets into a single dataset instead of analyzing them independently, increasing the number of principal components included per dataset, and alternative clustering algorithms including graph-based clustering. These assessments are described in Supplementary Methods “Sensitivity analysis for model building”.

### Connecting new data with the existing databases

To demonstrate the ability to match datasets under new analysis to relevant published datasets, we applied RAVmodel on five TCGA datasets (Fig. 2A). Based on the correlation to principal components in these datasets, we identified RAVs specific to breast invasive carcinoma (RAV221 and RAV868) and to colon and rectal adenocarcinoma (RAV832). When RAVmodel was applied to the TCGA-BRCA dataset, RAV221 was assigned with the highest validation score (Fig. 2B, Supplementary Table 1) and the associated MeSH terms were mostly breast-related terms, such as ‘breast’ and ‘breast neoplasms’ (Fig. 2C, drawWordcloud function). We extracted three breast cancer studies contributing to RAV221 (Fig. 2D, findStudiesInCluster function). GSEA annotations on RAVs were queried and the top 10 enriched pathways were all breast-cancer associated (Fig. 2E, subsetEnrichedPathways function). We also checked RAV832 on its association with TCGA-COAD and TCGA-READ datasets. RAV832 was assigned with the second highest validation score for both COAD and READ datasets (Supplementary Fig. 4A and 4B, respectively) and contained MeSH terms such as ‘colorectal cancers’, ‘colon’, and ‘adenocarcinoma’ (Supplementary Fig. 4C). We also recognized that three out of five training data in RAV832 directly represented colon-associated illnesses (Supplementary Fig. 4D) and the top enriched gene set was an upregulated pathway in colorectal adenoma (Supplementary Fig. 4E). In summary, we confirmed that RAVmodel serves as a specific and robust index, coherently connecting expression profile, gene sets, related study studies and their associated metadata (Fig. 1B), ultimately enhancing the interpretation of new datasets in the context of existing databases.

**Fig 2.**
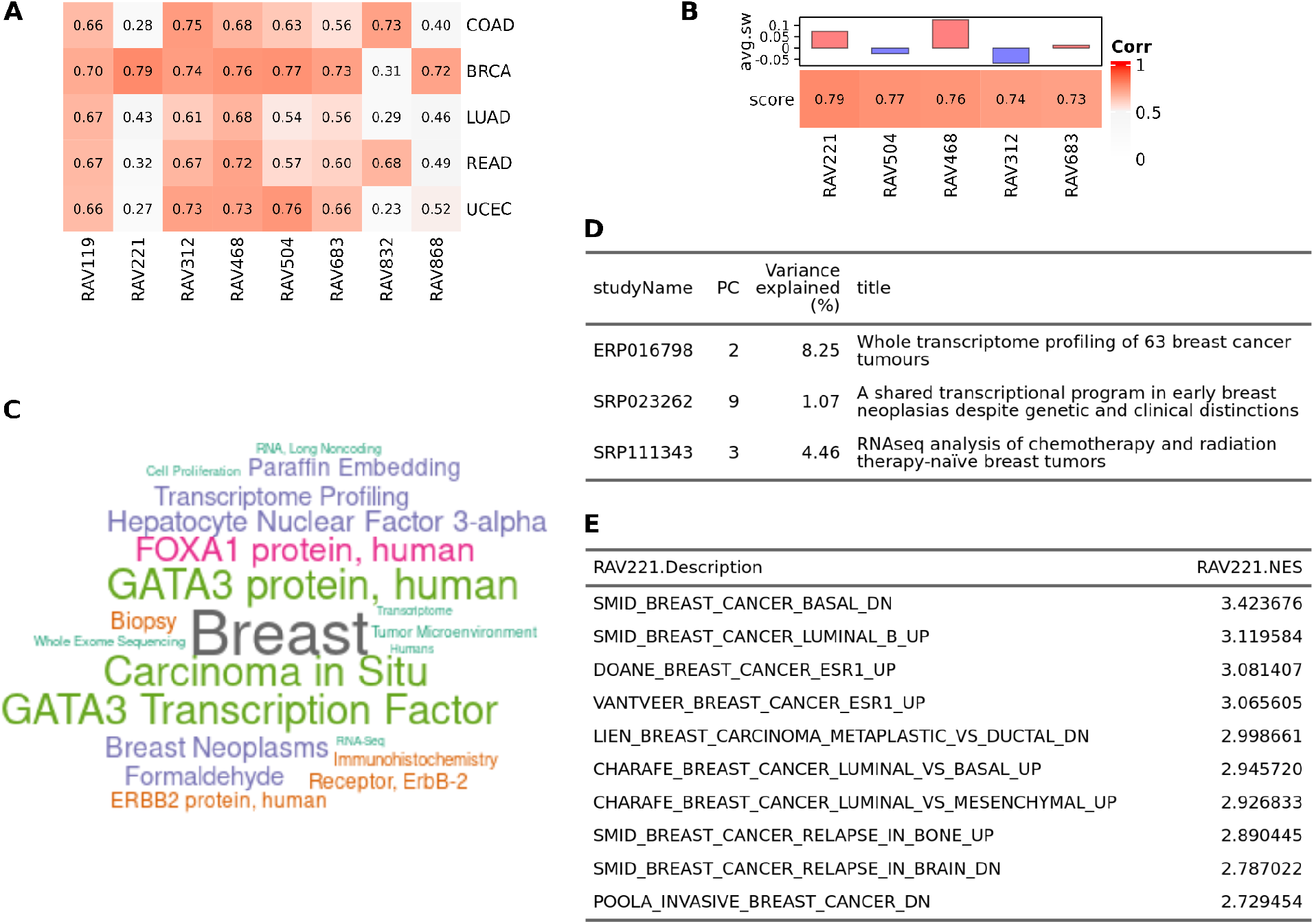
Connecting new datasets to existing databases. GenomicSuperSignature provides a rich resource for understanding new or user-supplied datasets in the context of existing datasets summarized in the RAVmodel. A) Validation of multiple TCGA RNA-seq datasets. Each dataset was subjected to PCA and Pearson correlation coefficients between top PCs and all possible RAVs were calculated. RAVs with Pearson coefficient above 0.7 in at least one dataset were displayed here. RAV221 and RAV868 indicate association with breast cancer while RAV832 is associated with colon and rectal cancer. (COAD: Colon Adenocarcinoma, BRCA: Breast Invasive Carcinoma, LUAD: Lung Adenocarcinoma, READ: Rectum Adenocarcinoma, UCEC: Uterine Corpus Endometrial Carcinoma) B) Validation of TCGA-BRCA. From panel A, we showed RAV221 is associated with breast cancer and confirmed RAV221 is one of the top validated RAVs for TCGA-BRCA. Top 5 validated RAVs (*score*, bottom panel) and their average silhouette width (*avg.sw*, top panel) are shown. C) A word cloud of MeSH terms associated with RAV221. We collected MeSH terms assigned to the publications belonging to RAV221 and weighted them based on their prevalence and the contribution to any given RAV (Supplementary Methods). This word cloud shows that RAV221 is heavily composed of principal components from studies of breast neoplasms. D) Three studies contributing to RAV221. E) Top 10 enriched pathways in RAV221.

### RAVs to characterize colorectal cancer

To compare the utility of GenomicSuperSignature relative to the focused use of data from a single disease, we compared RAVs to two previous studies that employed CRC gene expression databases to identify CRC molecular subtypes. The CRC Subtyping Consortium used 18 CRC datasets from multiple platforms comprising 4,151 patients to define four discrete Consensus Molecular Subtypes (CMS) observed across numerous patient cohorts^20,21^. Ma *et al.* subsequently proposed a continuous scoring system (PCSS) based on an analysis of 8 CRC microarray datasets comprising 1,867 samples, and found it was more closely correlated to microsatellite instability (MSI)^22,23^, grade, stage, and tumor location^20,21^. Importantly, these previous efforts both employed curated databases of only CRC transcriptomes, whereas the training set of the current RAVmodel consists of less than 2% CRC studies (Supplementary Table 3). We identified the RAVs most highly associated with CMS subtypes (RAV834/833) and PCSSs (RAV1575/834) (Supplementary Results), and confirmed that these RAV pairs showed comparable or higher performance on colon cancer subtyping than CRC subtyping efforts defined by bespoke methods in focused datasets (Fig. 3A, Supplementary Fig. 5A).

**Fig 3.**
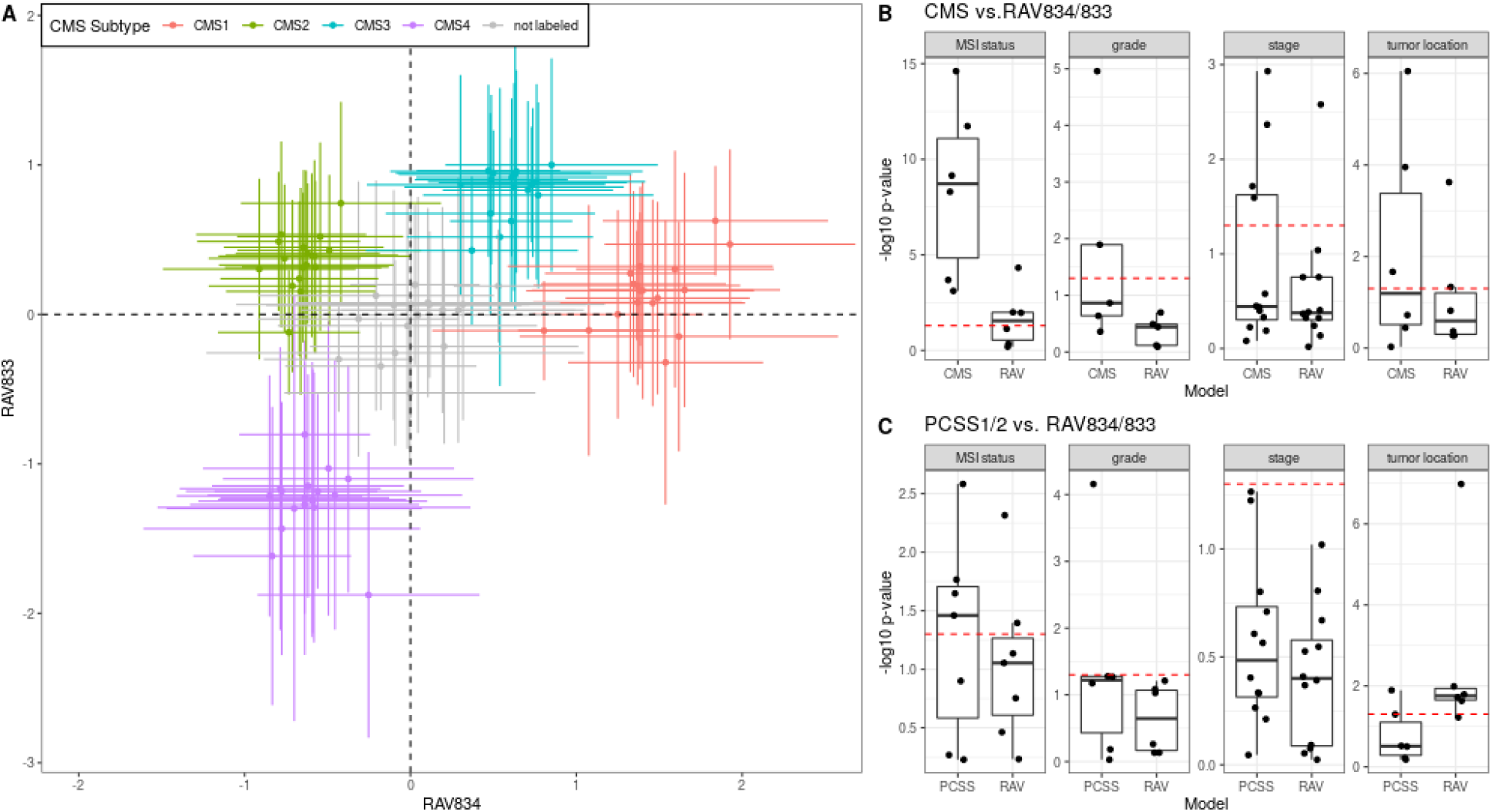
Sample scores for disease subtyping and metadata characterization. We assigned sample scores to 3,567 tumor samples from 18 CRC studies. A) The samples in each of 18 datasets, assigned to either (i) one of the 4 previously proposed CMS subtypes by CRC Subtyping Consortium^20,21^ or (ii) not assigned to a CMS subtype (so 5 × 18 = 90 total groups), are represented by the mean (point) and standard deviation (error bar) of sample scores. CMS subtypes (colors) separate when plotted in RAV coordinates. We further evaluated the capacity of RAVs to demonstrate clinicopathological characteristics of colon cancer. B) Clinical phenotypes were regressed on discrete CMS subtypes and RAV834/833-assigned sample scores as covariates. LRTs were used to compare the full model to a simplified model containing only CMS subtype (left) or RAV834/833-assigned sample scores (right) as predictor. RAV834/833-only model shows −log_10_p-value near 0, implying that CMS is not providing additional information. C) The same regression and LRTs as in panel B were done using PCSS1/2 and RAV834/833-assigned sample scores as covariates. RAV834/833 outperforms PCSS1/2 on explaining colon cancer phenotypes except tumor location.

Using training and validation data of the original CRC studies, we compared associations between different subtype models and RAVs with the same clinicopathological variables. Notably, these data were *not* part of RAV training and are microarray datasets whereas the RAVs were trained exclusively from RNA-seq data. As described previously^20^, we used the likelihood-ratio test (LRT) to compare the different subtype models for association with clinicopathological variables. A p-value near 1 (−log_10_p-value near 0) means that no additional information is provided by a full model composed of two subtype definitions compared to a model with only one. CMS-associated RAVs performed better than discrete CMS on all four phenotypes and also outperformed PCSSs except on tumor location (Fig. 3B-C). Interestingly, PCSS-associated RAVs were still better than CMS but slightly worse than PCSSs, while CMS-associated RAVs were better than both CMS and PCSSs, indicating that RAVs contain more comprehensive information (Supplementary Fig. 5B-C). This performance improvement became more significant using only the 10 original validation datasets, excluding 8 datasets used to train the PCSS model (Supplementary Fig. 6). In conclusion, RAVs trained from heterogeneous datasets, not specific to CRC, captured biologically-relevant signatures for CRC as well or superior to focused efforts using CRC-specific databases, suggesting that RAVs are for general-use and can be applied to describe other diseases as well.

### Identify common biological attributes across different datasets

For practical and technical reasons, biological datasets often contain missing information or signals buried in noise. GenomicSuperSignature can fill out those gaps by uncovering weak or indirectly measured biological attributes of a new dataset by leveraging the existing databases. To evaluate this ‘transfer learning’ aspect of the GenomicSuperSignature, we compared the neutrophil count estimation by RAVs across two different datasets, systemic lupus erythematosus whole blood (SLE-WB)^24^ and nasal brushing (NARES)^25^ datasets as described in the previous study^8^. We searched for the SLE pathology-relevant RAV in three different ways using the SLE dataset^24^. First, we identified RAV1551 based on the highest validation score with the positive average silhouette width. Second, we searched for the keyword, *neutrophil*, in GSEA-based annotation. Eleven RAVs, including RAV1551, had two keyword-containing, enriched pathways. Lastly, we used the neutrophil count of the SLE-WB dataset to find the metadata-associated RAV. For the continuous variables like neutrophil count, we compared the R^2^ between the target variable and all RAVs, where RAV1551 showed the maximum R^2^, 0.395 (Fig. 4A). Neutrophil is a terminally differentiated cell type and potentially under-detected in the active gene expression profile, so we used the neutrophil estimate from MCPcounter^26^ and further evaluated the correlation between RAV1551 score and neutrophil estimate as the previous study^8^. A stronger correlation between RAV1551 score and neutrophil estimate was observed (Fig. 4B). We concluded that RAV1551 is the SLE pathology-relevant RAV, specifically associated with neutrophil counts, and tested whether this information can be expanded beyond the SLE dataset. For that, we applied RAV1551 on NARES dataset, which is a gene expression profile of nasal brushings obtained from granulomatosis with polyangiitis (GPA) patients, a condition that causes inflammation of blood vessels affecting ears, noses, throats, lungs and kidneys^25^. RAV1551 was not a top validated signal, ranked 14th with the validation score 0.41 with PC1, implying that neutrophil phenotype is not a major feature of NARES dataset. However, R^2^ between the neutrophil estimate from MCPcounter and RAV1551 score was 0.84 (Fig. 4C). This suggests that RAV can serve as a new measure to compare different datasets and provide interpretation of potentially subtle biological signals.

**Fig 4.**
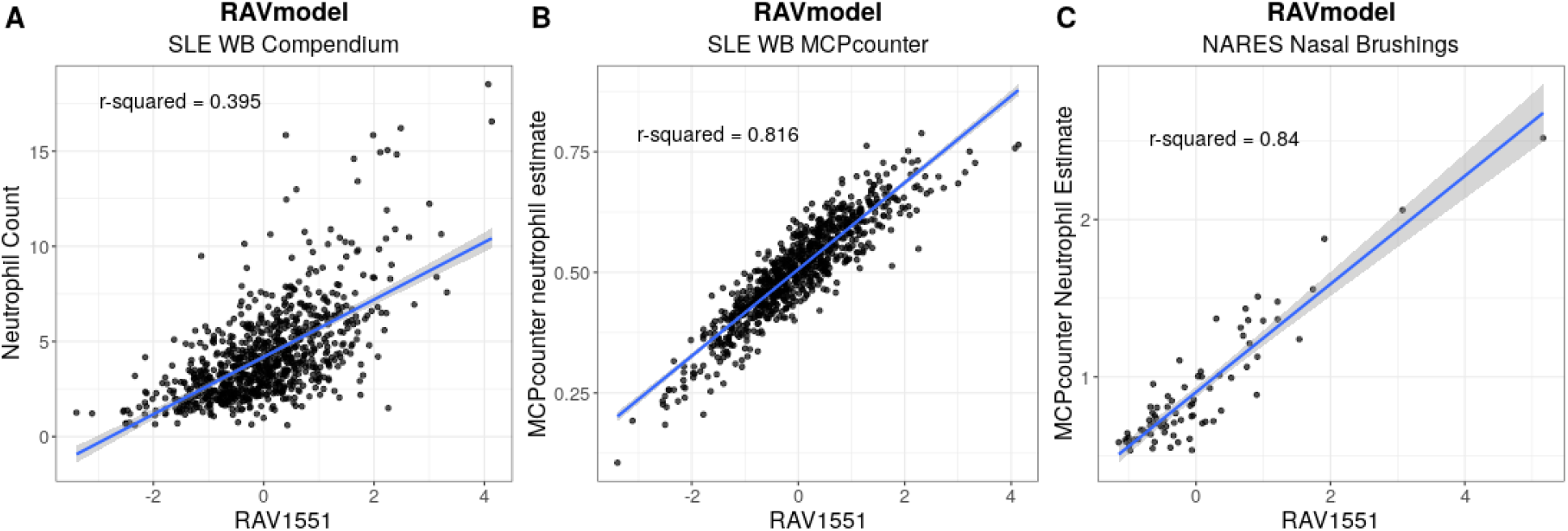
Estimate biological features of a new dataset using the signatures learned from public databases. RAVs encompass biological signals applicable across different platforms and independent datasets. We demonstrate this transfer learning capacity of RAVs by identifying the neutrophil-associated RAV from systemic lupus erythematosus whole blood (SLE-WB)^24^ data and using the same RAV to analyze nasal brushing (NARES)^25^ dataset. A) Neutrophil counts of SLE-WB data were plotted against RAV1551-assigned sample scores. B) Neutrophil count estimates by MCPcounter^8,26^ were plotted against sample scores assigned by RAV1551. C) Neutrophil count of NARES samples were estimated by MCPcounter and plotted against RAV1551-assigned sample scores. The shared area is the 95% confidence interval for predictions from a linear model.

## Discussion

A key innovation of GenomicSuperSignature is the creation of an index of ***r***eplicable ***a***xes of ***v***ariation (RAVindex) consisting of principal components repeatedly observed in independent analysis of multiple published datasets (Fig. 1A). Compared to approaches that merge training data, this strategy is highly scalable, can identify latent variables specific to small training datasets, and ignores technical artifacts that are not observed across multiple datasets. RAVindex is annotated with publication citations, *ME*dical *S*ubject *H*eadings (MeSH) terms, and gene sets, all of which is stored as the ‘RAVmodel’. Assembly of this information through RAVmodel creates an information resource which can be rapidly applied to new datasets on a standard laptop (Fig. 1B). GenomicSuperSignature augments standard RNA-seq exploratory data analysis by providing modes of interpretation and hypothesis testing that were previously impractical to apply.

GenomicSuperSignature contains information learned from a large body of existing studies that can be “transferred” to newly collected data. For example, the RAVindex contains cancer type-specific RAVs (Fig. 2A), including RAVs that are more closely related to colorectal carcinoma (CRC) clinicopathological variables than transcriptome subtypes previously identified through intensive analysis of CRC-specific databases bespoke subtyping efforts (Fig. 3, Supplementary Fig. 5). Such “transfer learning” is broadly applicable but particularly beneficial to the study of rare diseases and to small datasets where weak and under-represented, but biologically meaningful, signals can be identified^8^. To demonstrate this, we identified an RAV that was highly correlated to neutrophil content using a systemic lupus erythematosus (SLE) dataset not in model training data, and used this RAV to estimate neutrophil content in a nasal brushing (NARES) dataset that lacks neutrophil count information (Fig. 4). In addition to data inference, GenomicSuperSignature can be useful for analyzing disease progress, comparing phenotypes across independent datasets, and identifying weak biological signals. The current RAVindex contains 4,764 RAVs and 3,386 out of them are previously observed in two or more independent datasets, which can be expected to have many other such applications through transfer learning.

GenomicSuperSignature is expected to be robust to batch effects because it characterizes clusters of highly similar latent variables from two or more independent studies through cluster sizes, enabling to ignore any that are unique to a single study. We demonstrated the robustness of GenomicSuperSignature through sensitivity analysis and benchmarking against prior disease-specific analyses^8,20^. While trained exclusively on RNA-seq datasets, the performance of GenomicSuperSignature was not diminished when applied to microarray datasets.

Furthermore, we observed transfer learning functionality compatible with the results from recount2-MultiPLIER^8^, even though that model used a different matrix decomposition method, was trained at the sample level instead of dataset level, and used a training set with only ~10% overlap of samples (Supplementary Methods). We conclude that GenomicSuperSignature is robust to different technological platforms and to heterogeneity of training datasets, and enables interpretation of divergent datasets without subject-specific models.

GenomicSuperSignature offers significantly improved usability over the existing tools by adopting user-friendly application schemas. First, the pre-built models greatly reduce computational requirements for users: whereas training the current model took several days on 24 cores with 128 Gb memory, its application can be performed in seconds on a conventional laptop computer. Its implementation as an R/Bioconductor package allows ready incorporation into widely-used RNA-seq analysis pipelines. GenomicSuperSignature R/Bioconductor package also enables a large research community to reuse public data for more accurate analyses of new data.

The approach taken for GenomicSuperSignature is flexible and can be extended to other large publicly available databases. We plan to develop RAVmodels trained on microarray, single-cell RNA sequencing, and spatial transcriptomic data, from model organisms such as mouse, and to metagenomic data from microbiome studies. Cross-species RAVmodels can help extend the discoveries from model organisms to humans^27^. These planned efforts will generate an expanded information resource with broader applicability and enhanced utility. For example, the GSEA annotation part of the model is independent from the RAVindex building process, so we can easily build multiple versions of RAVmodel with different gene sets or even any combination of gene sets. Also, we can connect RAVs with additional information on the training data because RAVs keep information on their specific source data. While the collection of RAVmodels grows as described, the GenomicSuperSignature package will be maintained as a stand-alone toolbox equally applicable to these different RAVmodels. GenomicSuperSignature and its associated data resources will provide biomedical researchers with a new set of data exploration tools exploiting knowledge gained from hundreds and eventually thousands of existing public datasets.

## Methods

### Source data

We used human RNA sequencing datasets from RNA-seq Sample Compendia in refine.bio^17^, which hosts uniformly processed gene expression data from EBI’s ArrayExpress, NCBI’s GEO and SRA. Data were downloaded on April 10th, 2020, and we selected studies based on the following criteria: 1) Exclude studies with more than 1,000 samples because they are more likely to be single-cell RNA sequencing datasets. 2) Exclude studies assigned with a *Me*dical *S*ubject *H*eadings (MeSH) term, “Single-Cell Analysis”. 3) Exclude studies with fewer than 50 successfully downloaded and imported samples (Supplementary Table 2 and Supplementary Fig. 2). Criteria 1 and 2 are not meant to entirely eliminate single-cell data but do serve to reduce the chance of including large, sparse datasets for which we plan to develop more specialized approaches. After filtering, the complete compendium includes 536 studies (defined as a single SRA study) comprising 44,890 samples.

### Processing training datasets

Training data included each sample’s `quant.sf` file from Salmon outputs^28^, not aggregated or normalized. We imported `quant.sf` files using `tximport`, scaling transcripts-per-million (TPM) using the average transcript length across samples and the library size (`“lengthScaledTPM”`)^29^, followed by the log2 transformation. Ensembl transcript names were converted into gene symbols using the AnnotationDbi package^30^. Row normalization was done on all samples together, not at the individual study level, because correcting variability at the study level could remove the signals we want to capture^31^. For model building, we used 13,934 common genes among 536 studies’ top 90% varying genes, where the variation cutoff was based on their study-level standard deviation (Supplementary Fig. 1A and Supplementary Methods).

### Build RAVmodels

We performed PCA on pre-processed gene expression matrices independently for each study using the `stats::prcomp` R function for each gene centered but not scaled. Loading vectors of top 20 PCs from 536 studies (total 10,720 PCs) were clustered via hierarchical clustering. For hierarchical clustering, we calculated the distances between loadings using Spearman’s correlation coefficient and clustered them with ward.D agglomeration method. The number of clusters was set to the minimum number that can separate up to 50 negative controls (Supplementary Methods and Supplementary Fig. 3). PCs in each cluster were averaged and the resulting ‘genes × averaged loadings’ matrix, RAVindex, was combined with associated metadata, GSEA and MeSH annotations into a unified data structure that we term a RAVmodel. We performed a detailed evaluation of model building criteria including varying gene sets used for annotation, the size of training datasets, the number of clusters, and so on (Supplementary Table 4, Supplementary Methods).

### Annotation

Gene Set Enrichment Analysis (GSEA) is a common approach used to supply biological interpretation to lists or sets of genes^32–34^ that has also been used to interpret biological signals in principal components^35^. We subjected each RAV to GSEA to aid in interpreting the biological signals associated with it. Genes were ordered by loading value from each RAV and supplied as a `geneList` input for `clusterProfiler::GSEA`^36^. We collected enriched pathways with Benjamini-Hochberg (BH) adjusted p-value < 0.05 and among them, the pathways with the minimum q-values. The subset of GSEA results - NES, `Description, pvalue`, and `qvalues` - were included in the RAVmodel. The version of RAVmodel used in this study is annotated with Molecular Signatures Database (MSigDB) curated gene sets (C2, version 7.1)^32,37^, excluding any MSigDB C2 gene set with fewer than 10 genes or more than 500 genes.

MeSH terms^38^ were assigned to each study using the NCBI Medical Text Indexer (MTI) tool^39^. The relevance of MeSH terms in each RAV was assessed through the bag-of-words model: all the MeSH terms associated with the training datasets were considered as the ‘universe’ and each term in the cluster was reverse-weighted by the frequency of the given term in the universe. MeSH terms were also weighted by the variance explained by the principal component that they came from (Supplementary Methods).

### Validation datasets

Five TCGA RNA sequencing datasets (COAD, BRCA, LUAD, READ, and UCEC) were acquired from GSEABenchmarkeR^34^. Any genes with count-per-million (CPM) less than 2 were excluded and the count matrix was log2-transformed and centered but not scaled before PCA. Eighteen colon cancer microarray datasets from curatedCRCData were also used for validation^40^. Missing and infinite values were removed from these microarray data and the remaining expression values were centered at each gene level. To evaluate the ability to perform transfer learning to metadata, we used preprocessed versions of nasal brushing (NARES)^25^ and the systemic lupus erythematosus (SLE) datasets^24^.

## Data Availability

The datasets analyzed during the current study can be found at https://doi.org/10.6084/m9.figshare.19087691.v1. Training datasets used for the current RAVmodel are available at refine.bio RNA-seq sample compendia (https://www.refine.bio/compendia?c=rna-seq-sample).

## Code Availability

The workflow to build the RAVmodel is available from https://github.com/shbrief/model_building. All analyses presented here are reproducible using code accessible from https://github.com/shbrief/GenomicSuperSignaturePaper/. GenomicSuperSignature package is available at https://doi.org/doi:10.18129/B9.bioc.GenomicSuperSignature.

## Competing interests

The authors declare the following competing interests: Daniel Blankenberg has a significant financial interest in GalaxyWorks, a company that may have a commercial interest in the results of this research and technology. This potential conflict of interest has been reviewed and is managed by the Cleveland Clinic.

## Supplementary Materials

### Supplementary Methods

#### Sensitivity analysis for model building

##### Datasets for methods searching

During the optimization process for RAVmodel building, we used the well-characterized and smaller datasets: 8 colon cancer datasets from curatedCRCData, 10 ovarian cancer datasets from curatedOvarianData, and recount2 datasets used for recount2-multiPLIER model. Training datasets for recount2-multiPLIER model and the current version of RAVmodel are partially overlapping: recount2-multiPLIER model used 37,027 runs from 30,301 unique samples from 1,466 studies and GenomicSuperSignature was constructed from 44,890 runs from 34,616 unique samples from 536 studies. Among them, only 6,839 runs from 5,260 unique samples from 87 studies were used by both models. In addition to the different combinations of these datasets, we created synthetic datasets that can serve as positive and negative controls.

##### Dimensionality reduction methods

We assessed multiple dimensionality reduction methods for RAVindex building. Nonnegative matrix factorization (NMF) was excluded because there is no clear criterion to select representative components, such as variance explained by each principal component in PCA. Non-orthogonal relationship between components captured by NMF is potentially a more relevant representation of biological data, but by combining replicative principal components, we overcome the orthogonality constraint imposed by PCA. We also ruled out independent component analysis (ICA) because it separates independent signals to reduce the effect of noise or artifacts^41^, which is different from our goal to extract biological signals, and does not rank its components like NMF. We narrowed down to two dimensionality reduction methods, PCA and PLIER^7^, and investigated the types of signatures when they were applied at the dataset-level or sample-level. This comparison was done within four different conditions: perPCA (PCA on each dataset and cluster top PCs), megaPCA (PCA on all samples), perPLIER (PLIER on each dataset and cluster latent variables (LVs, equivalent to principal components from PCA)), and megaPLIER (PLIER on all samples, identical to multiPLIER). One of the downsides in megaPLIER approach was that the direct link between LVs and the training data was not available. Also, the annotation database was inseparable from the model building, making it harder to scale. perPLIER approach blended LVs in each cluster and lost distinct signatures. Like megaPLIER, megaPCA did not maintain the links between signature and it’s source data. Additionally, megaPCA picked up only a handful of strong signatures in top PCs, which we can still capture through the perPCA approach. Overall, we decided to use the perPCA approach for our model building because it is more scalable, keeps the link between signature and source data, and captures both pan-data and per-data signatures.

##### Data transformation

We did log2-transformation and row-normalization across all samples, not at the dataset level, to keep the differences in the datasets. VST transformation was excluded because it requires significantly more computing resources - over 200 times longer user CPU times, without any meaningful improvement on capturing biological signatures over log2-transformation because we removed low variable genes from our training datasets^42^.

##### Subset genes

We searched for the minimum set of genes that keeps the biological information, because more genes require more computing resources to process and some genes were measured only in certain training datasets. Also, low- or non-expressing genes can be similar to the background noise, so including them would make interpretation harder. First, instead of using a fixed cutoff for ‘low-expressing’ genes, we selected genes based on their expression variance within the dataset because we suspect that genes with the stable expression level within a dataset convey less information to capture. We also tried the common genes between training datasets and annotation databases before PCA. However, it didn’t improve the accuracy of GSEA compared to the common genes among training datasets and made the model less scalable because RAVindex needs to be rebuilt for RAVmodels with different annotation databases even for the same training datasets. So for the model building, we used the common genes among the top 90% varying genes from each training dataset.

##### The number of PCs to collect

We decided on the number of PCs to collect based on the following four reasons. First, a threshold of 20 PCs is adequate for stability of the RAV model, particularly for larger clusters of 3+ PCs. Most of the lower PCs (PC11-20) are in single-element (22%) or two-element (53%) clusters. We expect further relaxing the cutoff would contribute even less to clusters of 2+ size, which we validate using two RAVmodels consisting of 1) top 10 PCs (RAVmodel_10) or 2) top 20 PCs (RAVmodel_20) from each dataset. We chose the most similar pairs of RAVs between 2,382 RAVs (RAVmodel_10) and 4,764 RAVs (RAVmodel_20) using the Pearson coefficient. 79% of RAVs in RAVmodel_10 have a similar or identical RAVs in RAVmodel_20 with the Pearson coefficient > 0.7 and the average Pearson coefficient for the RAVs with more than 2 elements is 0.86, suggesting that the model is robust to cutoffs of >= 10 PCs, and that the added computational cost of a cutoff larger than 20 would provide little or no benefit. Second, we chose the top 20 PCs because they represent a majority of the gene expression variance in each study (median percentage of total variability represented 63%). Third, we applied the elbow method to find the number of ‘significant’ PCs to collect, using the `num.pc` function implemented in the PLIER package with the following modifications^18^. The `PLIER::num.pc` function applies z-score normalization, but because our method does normalization with all samples combined, we removed this internal normalization from `PLIER::num.pc` and provided the pre-normalized input data instead. The number for significant PCs from this modified `PLIER::num.pc` function ranged from 5 to 45, while the “elbow” of the scree plots was not always clear on manual inspection. We chose the median value, 20, as a pre-set cutoff for the different training datasets. Using a varying number of PCs would add a complexity to the process that seemed unjustified given that the variance explained by each PC does not vary much by study size, ranging from 50 to 100 for our 536 training datasets. For example, after the 8th PC, less than 5% of the variance was explained by a single PC for all 536 training datasets - the maximum variance explained by PC7 and PC8 are 5.1% and 4.6%, respectively. Finally, one of the main works we benchmarked against was Ma et al., where the authors selected top 20 PCs for their model building to extract colon cancer specific signatures.

##### Clustering methods

To group the replicative PCs, we tried centroid-based clustering such as k-means, graph-based clustering, and connectivity-based clustering like hierarchical clustering. We tried them on top 20 PCs from 8 colon cancer datasets and for evaluation, the cluster membership was compared with previously identified signatures (PCSSs) using Jaccard index. We also tested them on top 5 PCs from 10 positive and 10 negative controls, which were synthetic datasets created through bootstrap and random sample selection, respectively. We evaluated each clustering method based on how well top PCs from positive controls were clustered together and top PCs from negative controls were assigned into different clusters.

For the centroid-based clustering methods (k-means and k-medoids), we searched the optimum number of clusters using multiple measures including elbow method, mean silhouette width, and within sum of squares. However, the number of clusters required to separate unrelated PCs was too high to keep the related PCs together, which couldn’t be improved with different distance metrics. We suspect that PCs from biological data do not possess spherical or ellipsoidal symmetry required for centroid-based clustering to work.

We evaluated graph-based clustering and hierarchical clustering with the different combinations of distance metrics and agglomeration methods (for hierarchical clustering) on the same datasets used for the centroid-based clustering tests. From this evaluation, the clustering schema was narrowed down into two versions: for graph-based clustering, edge-betweenness clustering using edge weight by Spearman correlation and for hierarchical clustering, using Spearman distance and ward.D agglomeration. When we applied these clustering approaches on larger datasets, however, there were scalability issues with graph-based clustering: it formed a very large cluster, containing more than 5% of PCs, that cannot connect related datasets due to the extreme distribution of the cluster sizes. So we decided to use hierarchical clustering based on Spearman distance with ward.D agglomeration.

##### Choose the optimum number of clusters for hierarchical clustering

We used the negative-control dataset to explore the optimum number of clusters for hierarchical clustering. First, we constructed 50 synthetic datasets by randomly selecting 50 samples from 44,890 samples. We scrambled genes in each of 50 synthetic datasets and added random value between −0.1 and 0.1. The mean and standard deviation of 44,890 samples were used for row normalization of the synthetic datasets. We confirmed that these synthetics datasets can serve as a negative control based on the distance matrix: the minimum and maximum distance of PC1s from the synthetic datasets ranged approximately between 1st and 3rd quarters of the distance distribution of PCs from the actual training datasets, which we want to separate during the clustering process (Supplementary Fig. 3A). We collected top 20 PCs from 536 training datasets and PC1s from varying numbers of synthetic datasets (10, 20, 30, 40, and 50) and performed hierarchical clustering with the different numbers of clusters. Nine different cluster numbers were applied to each datasets: those nine cluster numbers were `round({#ofPCs}/d)`, where d is 7, 6, 5, 4, 3, 2.75, 2.5, 2.25, and 2. All negative controls were separated when `d = 2.25`, regardless of the number of negative controls (Supplementary Fig. 3B and 3C). So for the current versions of RAVmodel, we selected 4,764 clusters (= `round((20×536)/2.25)`).

##### Model validity

To test whether heterogeneous datasets can maintain the signatures from the focused dataset, we first built RAVindex from the focused training datasets and gradually diversed the training datasets with the unrelated datasets. A rate of overlapping enriched pathways over correlated pathways was monitored, from which we confirmed that our RAVindex building process reliably maintains the dataset-specific signatures from the heterogeneous training datasets.

#### Weighting MeSH terms in each cluster

The significance of MeSH terms associated with each cluster was evaluated based on their exclusivity. However, the simple sum of associated MeSH terms can be inappropriate in some cases. For example, potential noises can be a predominant signal in small clusters and common MeSH terms, such as ‘human’ or ‘RNA sequencing’ for the current version of RAVmodel, can be overrepresented and cover all the other terms. To handle these extreme situations, we applied several filtering and normalization terms. If the cluster contains less than 8 PCs, we considered any MeSH terms appearing half down of ‘cluster size x 0.5’ as noise and removed them. If the cluster has more or equal to 8 PCs in it, any MeSH terms appearing less than or equal to 4 times were eliminated. These cutoff values for ‘noise’ can be modified by users to fit their needs. Remaining MeSH terms were assigned with the new score based on the variance explained by PCs and the sum of scores for a given MeSH term was divided by the frequency of that term in the universe, where the ‘universe’ is defined as the complete pool of MeSH terms associated with all the training datasets. This final score can be displayed as a table or a word cloud, using `meshTable or drawWordcloud` functions, respectively.

#### Prepare input dataset

The GenomicSuperSignature takes gene expression profiles generated from both microarray and RNA sequencing with the minimum pre-processing. The major requirement is that gene expression profile should follow the normal distribution.

### Supplementary Results

#### Software implementation

The PCAGenomicSignatures class inherits SummarizedExperiment data structure and stores RAVindex, metadata, and annotation, which we collectively refer to as the RAVmodel (Supplementary Fig. 1B). Functions and S4 methods for the PCAGenomicSignatures class to access components of the RAVmodel, visualize analyses, and interpret new datasets are implemented in the GenomicSuperSignature R/Bioconductor package (Supplementary Table 1). We provide different versions of RAVmodels based on gene sets used for GSEA annotation and they are readily available (as .rds format files) to download from the internet through the getModel function or wget (Supplementary Table 4).

Two key functions, `validate` and `calculateScore`, allow interpretation of new datasets at the study level and at the individual sample level, respectively. The validate function calculates Pearson correlation coefficients between the top 8 PCs of a dataset and all RAVs (Supplementary Methods), from which the highest value is assigned as a ‘validation score’ of the corresponding RAV. Validation score provides a quantitative representation of the relevance between a new dataset and RAV. In general, the higher validation score implies that the RAV explains a more significant feature of a dataset. Validation outputs can be visualized as a heatmap table (Fig. 2A and 2B) and an interactive plot (Supplementary Fig. 7), through heatmapTable and plotValidate functions, respectively. Average silhouette width of each cluster is available as a reference for quality control and as an additional filtering option to find significant RAVs. The calculateScore function calculates a RAV-assigned ‘sample score’ to each sample, which is the matrix multiplication result (B^t^, red) between the ‘samples × genes’ matrix (Y^t^, grey) and RAVindex (Z, blue) (Supplementary Fig. 1C). Similar to validation score, sample score provides a quantitative representation of the relevance between samples and the given RAV.

In addition to the study-level validation scores and the sample scores acquired through gene expression profile, we can access the knowledge comprising GenomicSuperSignature through various entry points, such as metadata, MeSH term, and keywords, because RAVmodel maintains the link between RAV and it’s source data. (Fig. 5).

#### Comparison to existing tools

We compared GenomicSuperSignature with the two existing methods using large databases for transfer learning, MultiPLIER^8^ and weighted nearest neighbor (WNN)^14^ (Supplementary Table 7 and Supplementary Table 8). MultiPLIER is a transfer learning framework for rare disease study and its signal is named as latent variables (LVs) which are similar in concept to RAVs. We included a biological example inspired by MultiPLIER (Figure 4). Our approach, while arriving at a similar result for this example, differs fundamentally from the approach applied in the MultiPLIER and offers distinct advantages. Whereas MultiPLIER identified the neutrophil-associated signal with the help of a relevant pre-established, single-dataset PLIER model, our method recovers this neutrophil signal in three unsupervised steps: validation of dataset signals through matching to RAVs, keyword searching against enriched gene sets for RAVs, and neutrophil-associated metadata.

With respect to implementation, GenomicSuperSignature software and RAVmodels differ from MultiPLIER in their construction process. In addition, RAVmodel provides versatile functionalities including software tooling for directly indexing new data against the RAVs and annotation of principal components of new data. Below we list some of the novel aspects and differences between the RAVmodel and the MultiPLIER method and its implementation.

1. MultiPLIER uses a dimensional reduction method called PLIER. RAVmodel uses principal component analysis followed by clustering of similar PCs from large training datasets.
2. For MultiPLIER, the PLIER dimensional reduction process is constrained by gene sets, so different gene sets require independent models even though the training datasets are identical. However, because gene set annotation of RAVmodel is not a part of RAVindex building, a single RAVindex can be annotated with different gene sets. This modularity and flexibility is one of the strengths RAVmodel has over the existing tools.
3. Unlike LVs, RAVs maintain the information on which primary studies contribute to the signal: the publication, the PC, and the variance explained by the PC. This metadata enables the direct connection of new data to individual existing studies. As a result, any new knowledge and metadata associated with the training datasets can be immediately incorporated into the RAVmodel and used for new data analysis.
4. Software: RAVmodel is a component of our GenomicSuperSignature Bioconductor package and Galaxy web tool for easy application of the model on new data. However, MultiPLIER is solely a transfer learning model and no specific software is provided for its application.
5. Model availability: To obtain the recount2 MultiPLIER model, users need to download a 81Gb zipped file from figshare, which includes 2.1Gb of model. On the other hand, different versions of RAVmodels are stored in Google Cloud bucket and users can download < 500Mb RAVmodels using `wget` or the `getModel` function provided in the GenomicSuperSignature package. The RAVmodels themselves are formalized as a subclass of the ubiquitous SummarizedExperiment Bioconductor class and, as such, will feel familiar to Bioconductor users.
6. Database search: RAVmodel enables unsupervised and coherent database search of sample metadata, study data, and PCs from user data, whereas MultiPLIER does not have any database search capability.
7. Enhanced interpretability of PCA: RAVmodel links PCs of the input data with the most relevant RAVs, which subsequently links PCs to the existing database of studies, sample metadata, and pathway enrichment. This process of ‘labeling PC’ is implemented as the `annotatePC` and `plotAnnotatedPC` functions in the GenomicSuperSignature package. There is no comparable functionality in MultiPLIER.

WNN is developed for transfer learning of single-cell multimodal data and implemented in Seurat^14^. For information transfer purposes, a reference atlas from 161,764 cells was built from 8 volunteers for HIV vaccine before and after the vaccine administration, which is not public data. These cells were thoroughly analyzed at the RNA and protein levels by the authors. Query data is linked to this reference via clustering (weighted-nearest neighbor) and the mutual nearest neighbor cells serve as ‘anchors’ to transfer information from the reference to query data. These algorithmic features and “training” data make Seurat’s WNN applicable largely to input datasets consisting of or containing PBMC for identification of immune cell type and states. The training data for RAVmodel is instead 44,890 bulk RNAseq data from 536 independent studies available through public archives. Instead of providing a reference map like Seurat’s WNN, our data compression procedure creates a data index, the RAVmodel, that connects literature, gene set annotation, sample metadata, and compressed gene expression signals. User-supplied, new query data can be linked to the RAVmodel through correlation between RAVs and query data’s PCs. RAV transfers information across different databases and independent studies in an unsupervised manner.

#### RAVmodel

RAVmodel is composed of RAVindex, model’s metadata, and annotation modules linked through RAVs (Supplementary Fig. 1B). RAVindex has 4,764 RAVs and 1,378 out of them are ‘single-element’ clusters. By definition a single-element cluster is not a ‘repetitive’ signal, leaving only 3,386 actual RAVs. This means we compressed the information from 44,890 samples into 3,386 RAVs, which is less than 1/10 of the initial number of samples. Also, 417 out of 536 training datasets have 40,746 genes and the other 119 training datasets have 41,255 genes, while the RAVindex uses only 13,934 common genes among the top 90% varying genes of all samples. Thus, our method achieves an efficient data compression, maintaining significant information in ~3% of the initial volume of the training data.

Cluster sizes, which is the number of PCs in RAVs, are strongly skewed to the right (Supplementary Fig. 9). When we exclude single-element clusters, about 65% of RAVs (2,212 out of 3,386) have two PCs and the mean cluster size is 2.759 PCs per RAV with the largest cluster containing 24 PCs. Interestingly, the distribution of PCs in RAVs changes with the cluster sizes: the majority of PCs in one- and two- element clusters are lower PCs, but once clusters have more than two elements, PC distribution starts to skew to the right (Supplementary Figure 10). This suggests that RAVs from small clusters tend to represent weak and less common signals. Though, we still keep the ‘single-element’ RAVs for two reasons: 1) If any new data is validated by those ‘single-element’ RAVs, they become ‘repetitive’ signals and thus, could lead to new hypotheses and 2) by keeping all RAVs, we include all potential PCs in the RAVmodel and support different use cases. Since metadata associated with all RAVs are readily accessible, end users can filter downstream results based on cluster sizes or other properties.

We assess the number of enriched gene sets for each RAV. About 40% of RAVs in RAVmodel_C2 and 50% of RAVs in RAVmodel_PLIERpriors do not have any enriched pathway and the majority of them are one- or two- element clusters (Supplementary Fig. 11), suggesting that the smaller clusters are less likely to represent biological features. Because there are RAVs annotated with only one input annotation, MSigDB C2 or PLIERpriors, we include all the RAVs to make our model cover diverse annotation databases. We further evaluate the scope of biological features represented by RAVmodel through two model validation measures, pathway coverage and pathway separation, adopted from the previous study^7,8^. Pathway coverage is defined as the proportion of pathways annotating RAVs out of all the gene set terms provided. Pathway coverage of RAVmodel_C2 is 0.32. The recount2_MultiPLIER has the pathway coverage of 0.42 while the RAVmodel_PLIErpriors which uses the same gene set as recount2_MultiPLIER has 0.64 pathway coverage. Pathway separation is defined as the ability of the signature model to keep non-overlapping signatures that can differentiate biologically similar pathways. Three biological subjects were tested on RAVmodel_PLIERpriors - type I versus type II interferon, neutrophil versus monocyte, and G1 versus G2 cell cycle phases. RAVmodel can successfully separate them either with top one or top five enriched pathways.

Redundancy within the cluster is defined as the cluster containing more than one PCs from the same study. The majority of RAVs (78%, 2,628 out of 3,386 non-single-element RAVs) consist of PCs from unique studies. 622 non-single-element RAVs are composed of only one study and 80% of them have no or only one MSigDB C2 pathway enriched.

To guide the interpretation, GenomicSuperSignature gives a message when the output included any of the following RAVs: 1) single-element RAVs, 2) RAVs with no or too many enriched pathways, where ‘too-many’ is defined as 5% of input gene sets (276 and 31 for MSigDB C2 and PLIERpriors, respectively), 3) non-single-element RAVs constructed from a single study. These criteria together include 2,557 RAVs.

#### Example of RAV interpretation

We identified a neutrophil-associated RAV, RAV1551, using biological information including enriched gene sets and validation of dataset by matching to RAVs. Here, we provide additional examples of RAV interpretation selected by the structure of RAVindex itself. As a first example, RAV3133 is the only single-element cluster consisting of a PC1, which is derived from SRA study SRP100652 containing 100 samples. This study investigated the gene expression effect of disease associated polymorphisms in the endoplasmic reticulum aminopeptidase genes ERAP1 and ERAP2^43^. PC1 of this dataset explains 16.2% of the variance and has only one enriched pathway with a very low NES. Interestingly, this dataset is zero-inflated (99.9% of counts are 0) and all the RAVs consisting of PCs from SRP100652 are tagged with the message implemented in GenomicSuperSignature as an interpretation guide. The other example is RAV2285 with a high proportion of PC1, where 15 out of 17 PCs in this RAV are PC1s while it also contains PC2 and PC5. Except one PC1, all the PCs in this RAV are from single cell RNA sequencing analysis (scRNAseq). The exception is PC1 from SRP116952, where RNA sequencing was performed on both total mRNA and polysome-associated mRNA^44^.

#### RAVs for colorectal cancer characterization

To evaluate the performance of RAVs compared to PCSSs, we searched for colon cancer associated RAVs in three different ways. First, we ran Kruskal-Wallis rank sum test between CMS subtypes and RAV-assigned scores. Two RAVs with the highest chi-square, RAV834 and RAV833, were selected for further testing. Second, we identified two RAVs, RAV1575 and RAV834, with the highest absolute Pearson correlation coefficients with PCSS1 and PCSS2, respectively (0.59 and 0.56). Last, we calculated validation scores for 18 colon cancer datasets from curatedCRCData^40^ and collected top 10 validated RAVs from each dataset. We summarized the frequency of different RAVs validating each dataset without any additional filtering criteria and selected the top 2 most frequently validated RAVs, RAV188 and RAV832, which were captured 14 and 10 times, respectively (Supplementary Table 5). In spite of the major difference in training datasets, RAV834/833 showed a comparable performance on colon cancer subtyping to PCSS1/2 (Fig. 3A). Notably, RAVs identified by CMS metadata (RAV834/833) performed better at CRC subtyping than the validated RAVs (RAV188/832), suggesting that the most prominent feature shared by 18 CRC datasets is not their disease subtypes (Fig.3A and Supplementary Fig. 5D).

#### PCA plot annotated with pre-calculated GSEA

One of the widely used exploratory data analysis methods is PCA and a PCA plot can provide a quick overview of sample composition and distribution. However, the interpretation of different PCs is not readily available in the conventional PCA. We couple PCs from new data with GSEA annotation of RAVmodel, and enable the instant interpretation of PCA results. We showed this example using a microarray dataset from isolated immune cells (E-MTAB-2452)^45^ and RAVmodel annotated with three priors from the PLIER package (RAVmodel_PLIERpriors, Supplementary Table 4). GenomicSuperSignature performs PCA on a centered, but not scaled, dataset and identifies the most relevant RAV for each PC. GSEA annotation of these matched RAVs can be summarized in a table (Supplementary Fig. 8A, `annotatePC` function). If you want to draw a PCA plot - currently any pair of top 8 PCs is supported, GSEA annotation will be displayed as a linked table (Supplementary Fig. 8B, `plotAnnotatedPCA` function).

## Supplementary Figures

**Sup.Fig 1.**
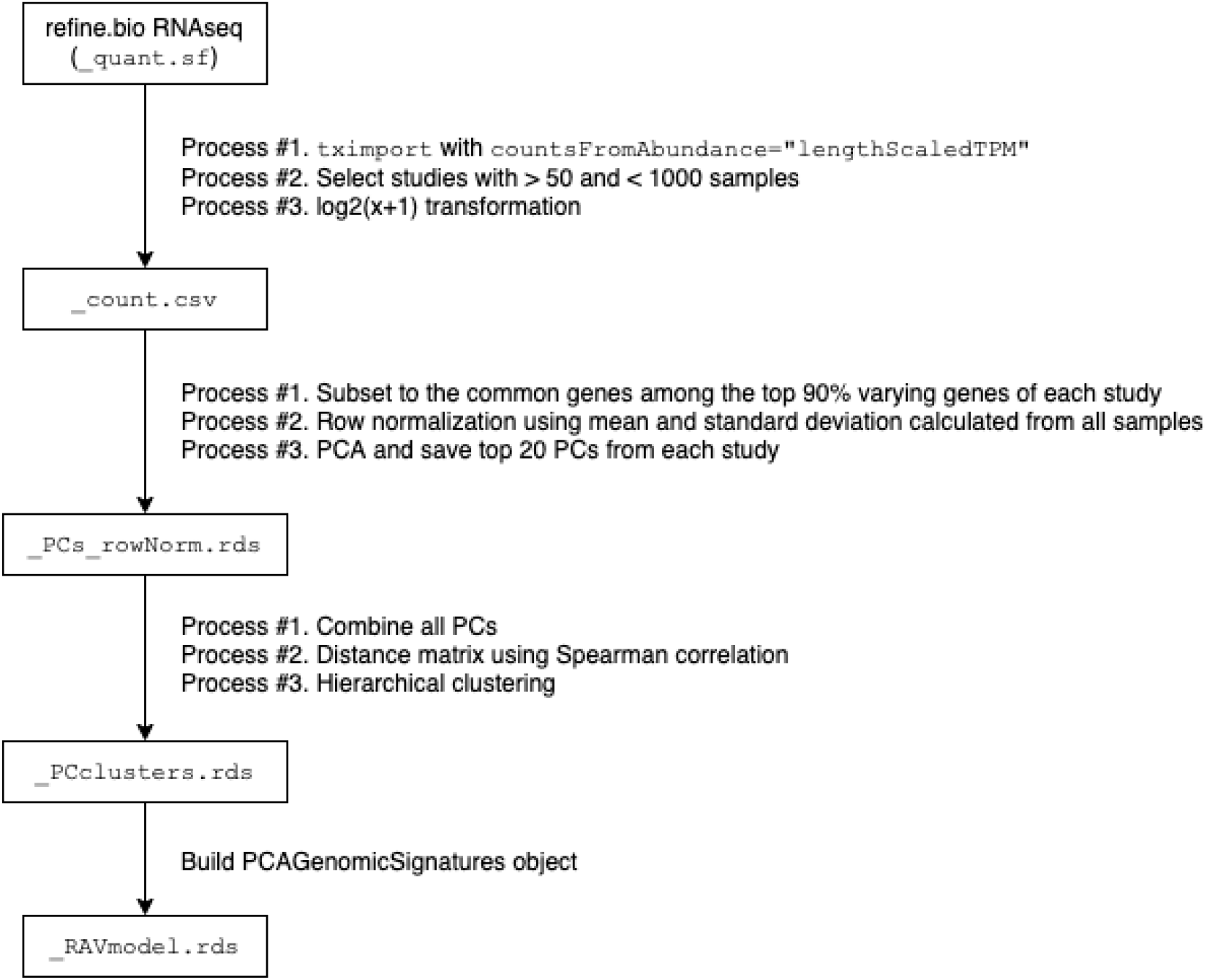
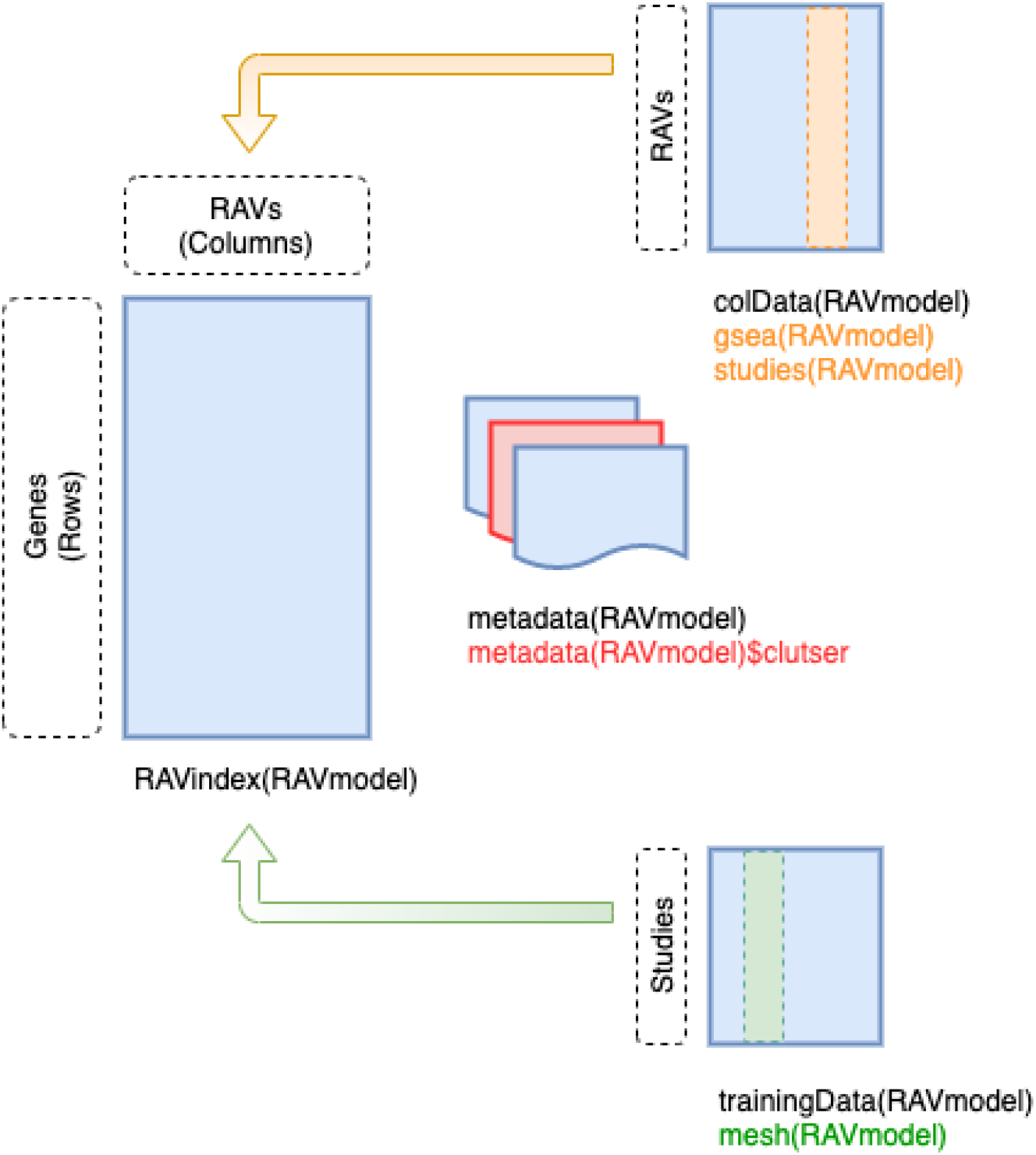
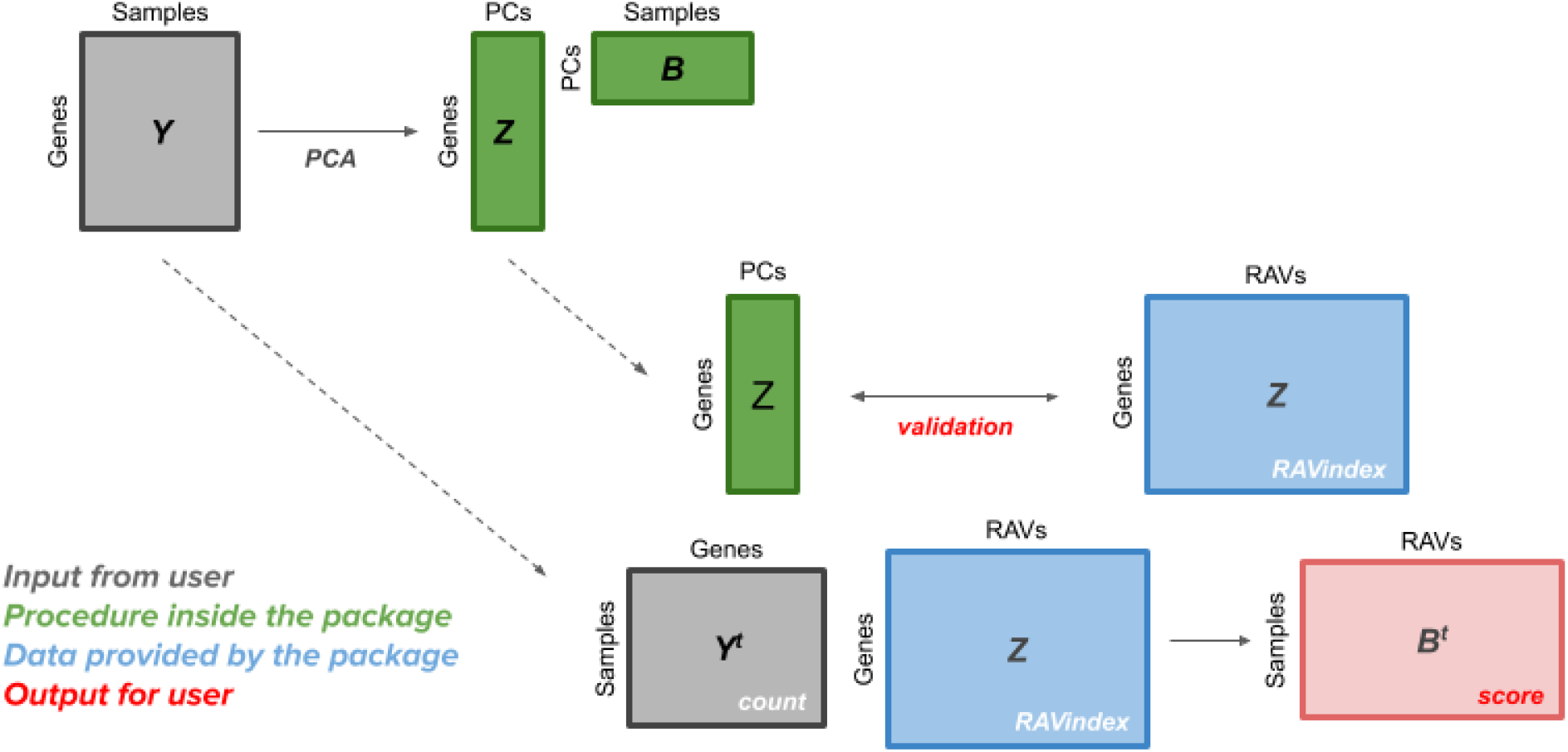
Overview of RAVmodel building. A) We downloaded human `quant.sf` files from refinebio RNA-seq Sample Compendia. We subset studies with more than 50 and less than 1,000 samples from 6,460 studies available at the time of the snapshot. Some RAVmodels retain additional filtering criteria on their training datasets. For example, the current version of RAVmodel predominantly used in this study further excludes datasets potentially from single cell analysis. Selected datasets were imported through `tximport`, followed by log2 transformation to bring them close to normal distribution. We used the common genes among the top 90% varying genes of each study, which was 13,934 genes for the current RAVmodel, and did row normalization using mean and standard deviation calculated from all 44,890 samples. PCA was done on a row-normalized expression matrix at the study level and top 20 PCs from each study were collected, ending up with 10,720 PCs. Distance matrix between these PCs was calculated using Spearman correlation and hierarchical clustering was applied with the pre-defined optimum number of clusters. Weighted MeSH terms and GSEA on each RAV, along with RAVindex and other metadata, were assembled into PCAGenomicSuperSignatures object, named as RAVmodel. In the below workflow diagram, boxes represent the intermediate files we created during the model building process. B) Schematic of PCAGenomicSuperSignatures object. RAVindex is a ‘genes × RAVs’ matrix. The colData provides information on RAVs, such as studies contributing to each RAV and GSEA results from each RAV. The metadata stores details on the RAVmodel itself, such as cluster memberships of PCs and the size of each cluster. The trainingData provides information on studies used for the model training, which includes MeSH terms assigned to each study and PCA summary of each study. C) User’s perspective. The GenomicSuperSignature package allows users to access a RAVmodel (Z matrix, blue) and annotation information on each RAV. From a gene expression matrix (Y matrix, grey), users can calculate dataset-level validation score or sample score matrix (B matrix, red). Through the RAV of your interest, additional information such as related studies, GSEA, and MeSH terms can be easily extracted.

**Sup.Fig 2.**
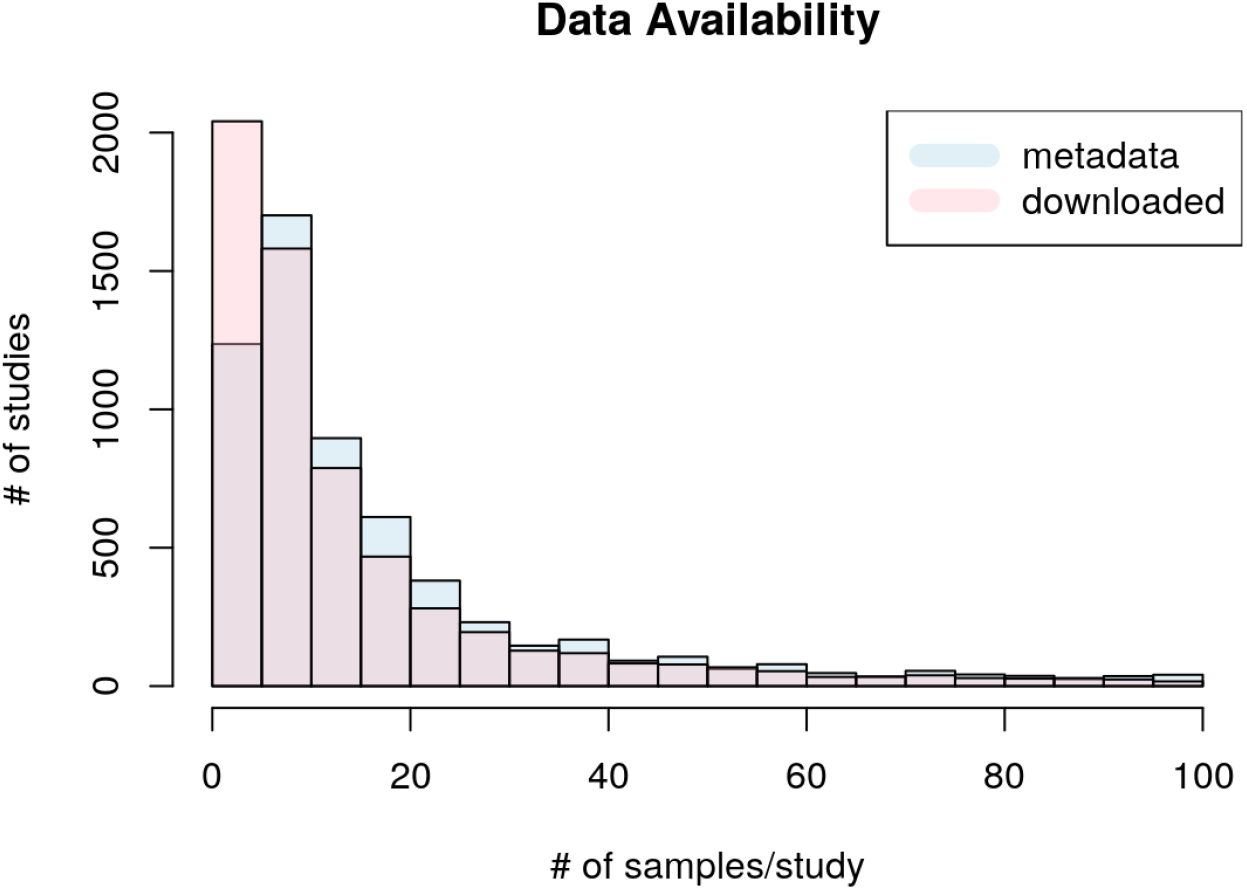
Data availability from refine.bio. The refinebio database is actively updated and our current RAVmodel is based on the snapshot on April 10th, 2020. Metadata bar (light blue) shows the number of studies with the given ranges of study sizes based on the metadata. Downloaded bar (pink) represents the number of studies with the given ranges of study sizes that were successfully downloaded and imported through `tximport`. Based on metadata, there were studies with more than 100 samples, but at the time of snapshot, only up to 100 samples were available. Thus, the plot displays only up to 100 samples/study cases. Due to the unavailability of certain samples, more studies belong to 0-5 samples/study bracket than metadata suggests.

**Sup.Fig 3.**
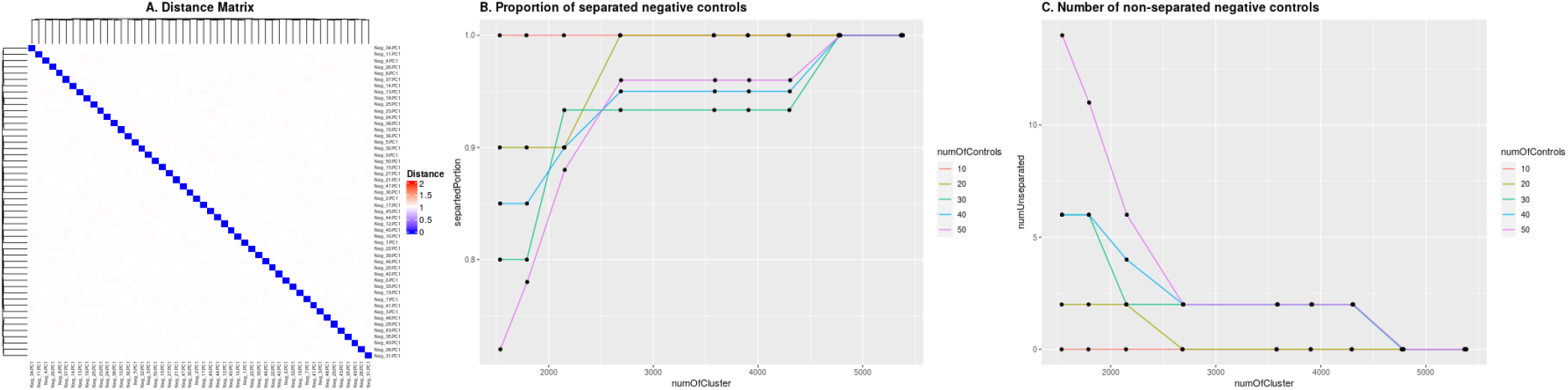
The optimum number of clusters. We used PC1s from synthetic datasets, designed as negative controls, to decide the number of clusters for hierarchical clustering. Training datasets, top 20 PCs from 536 studies (`RAVmodel_536` column in Supplementary Table 2), were combined with PC1s from the different numbers of synthetic datasets (10, 20, 30, 40, and 50) (Supplementary Methods). A) Heatmap of the distance matrix between 50 negative controls. Distance was calculated based on Spearman’s correlation. B) Proportion of the negative controls that were separated with the given cluster number. `numOfControls` is the number of negative controls added to the training datasets. `numOfCluster` is the round of the total PCs (from training datasets and negative controls) divided by 7, 6, 5, 4, 3, 2.75, 2.5, 2.25, and 2. Different numbers of negative controls were completely separated when we used the cluster number k = `round`((`the number of PCs)/2.25`). C) Number of negative controls that were not separated.

**Sup.Fig 4.**
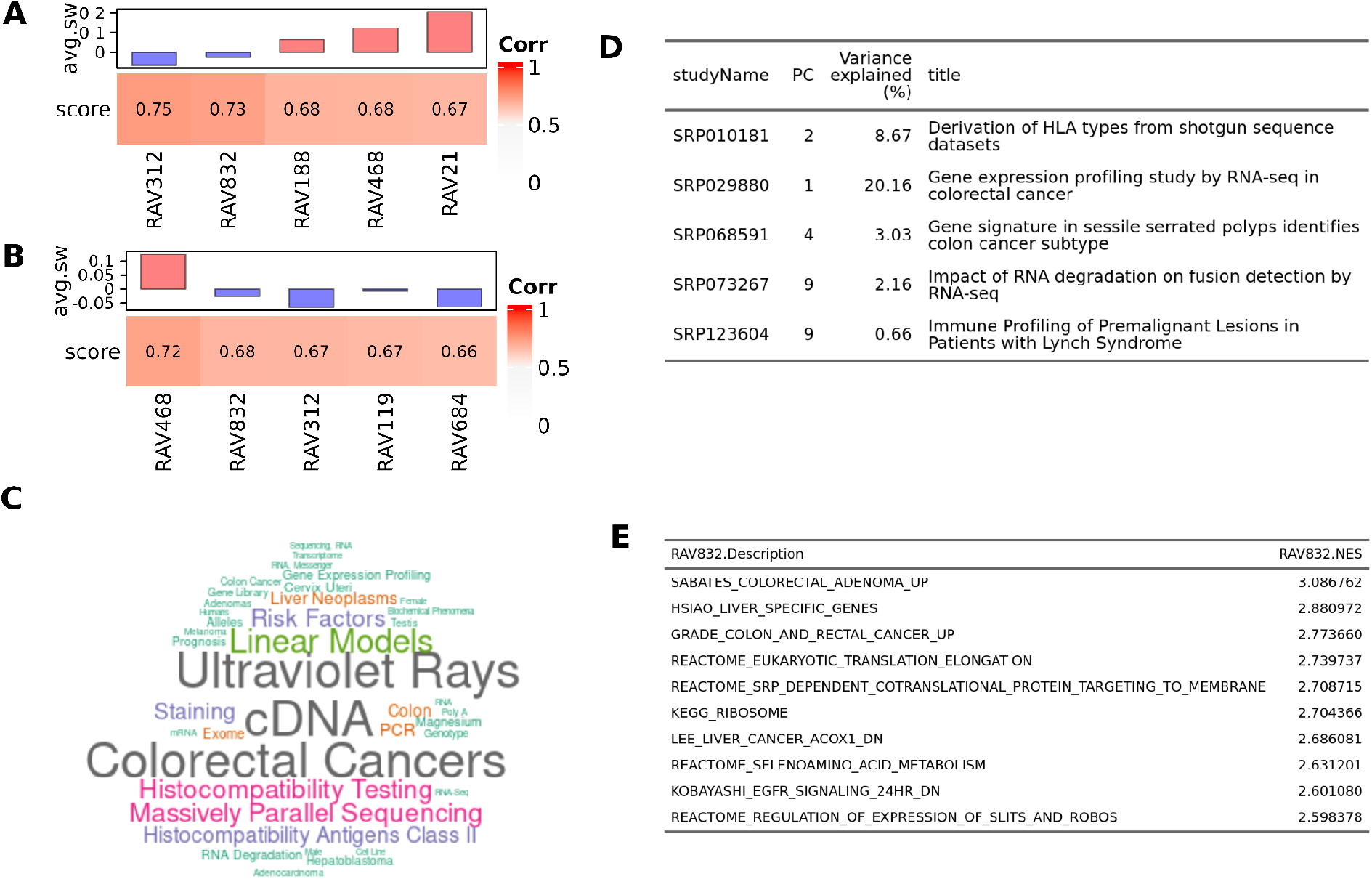
Colon and rectal cancer associated RAV. Based on Fig. 2A, RAV832 seems to be associated with TCGA-COAD and TCGA-READ. Top validation results of A) TCGA-COAD and B) TCGA-READ include RAV832 with the negative average silhouette width. C) MeSH terms associated with RAV832. D) Studies contributing to RAV832. E) MSigDB C2 gene sets enriched in RAV832.

**Sup.Fig 5.**
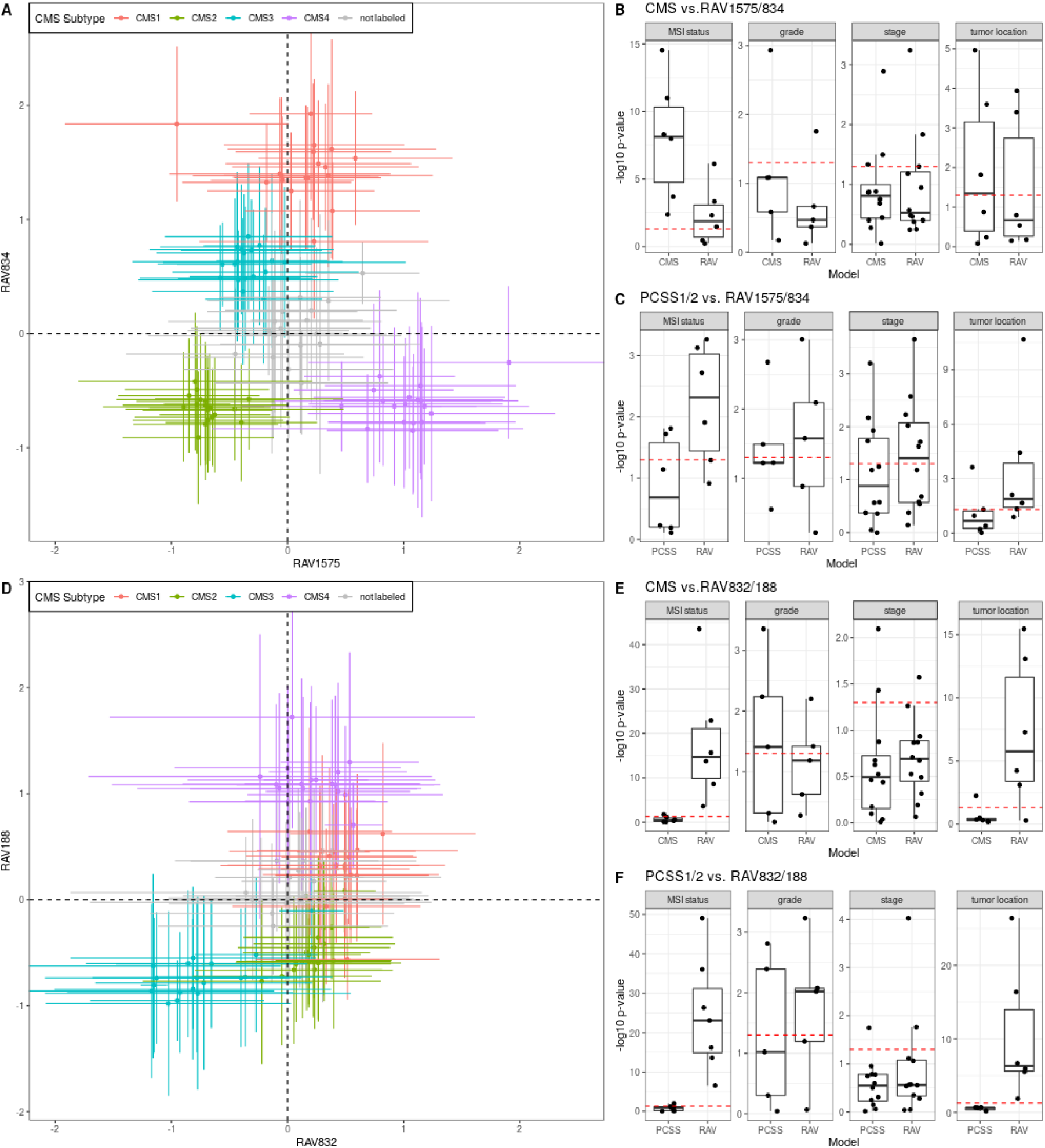
CRC characterization with different RAVs. In the results, we described two additional pairs of RAVs, RAV1575/834 and RAV188/832, that are potentially useful for CRC characterization. We applied the same analysis procedure on 18 CRC datasets as in Fig. 3 using those two pairs of RAVs. For the panel A and D, we assigned sample scores to 3,567 tumor samples from 18 CRC studies. The samples in each of 18 datasets, assigned to either (i) one of the 4 previously proposed CMS subtypes by CRC Subtyping Consortium^20,21^ or (ii) not assigned to a CMS subtype (so 5 × 18 = 90 total groups), are represented by the mean (point) and standard deviation (error bar) of sample scores. CMS subtypes (colors) separate when plotted in RAV coordinates. (A-C) CRC characterization with RAV1575/834. RAV1575 and RAV834 were identified based on their similarity to PCSS1 and PCSS2, respectively. (D-F) CRC characterization with RAV188/832. RAV188 and RAV832 were most frequently found among the top 10 validated RAVs.

**Sup.Fig 6.**
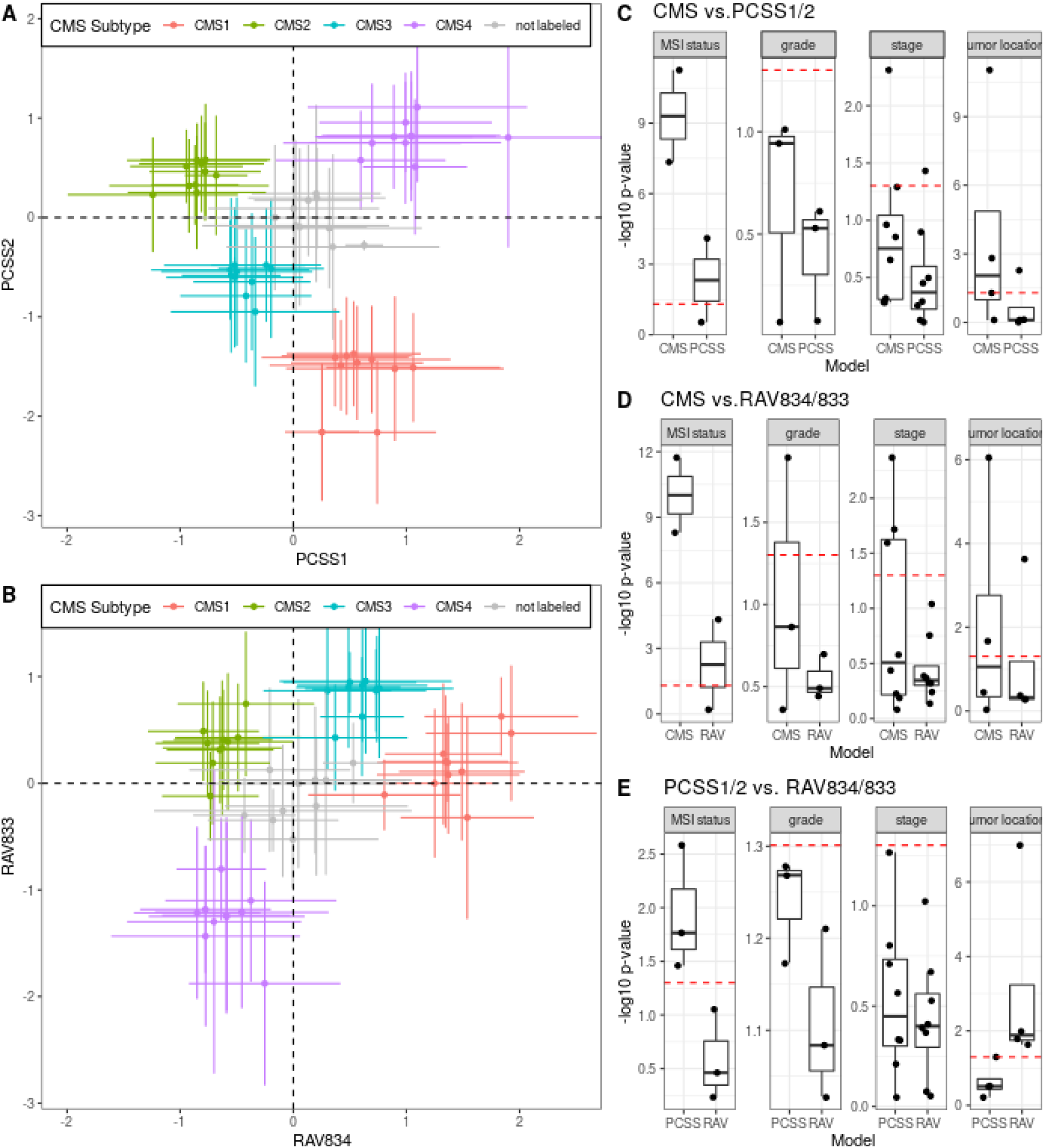
CRC characterization with 10 validation datasets. Analyses in Fig. 3 were repeated with only 10 CRC datasets, excluding 8 datasets used to train PCSSs. A) Subtype- and study-specific mean of PCSS1 and PCSS2 scores are plotted as points while the error bars represent standard deviation. B) The same plotting scheme as A was applied on RAV834 and RAV833-assigned scores. C-E) LRTs compare the full model to a simplified model containing only C) CMS subtypes or PCSS1/2, D) CMS subtypes or RAV834/833, and E) PCSS1/2 or RAV834/833.

**Sup.Fig 7.**
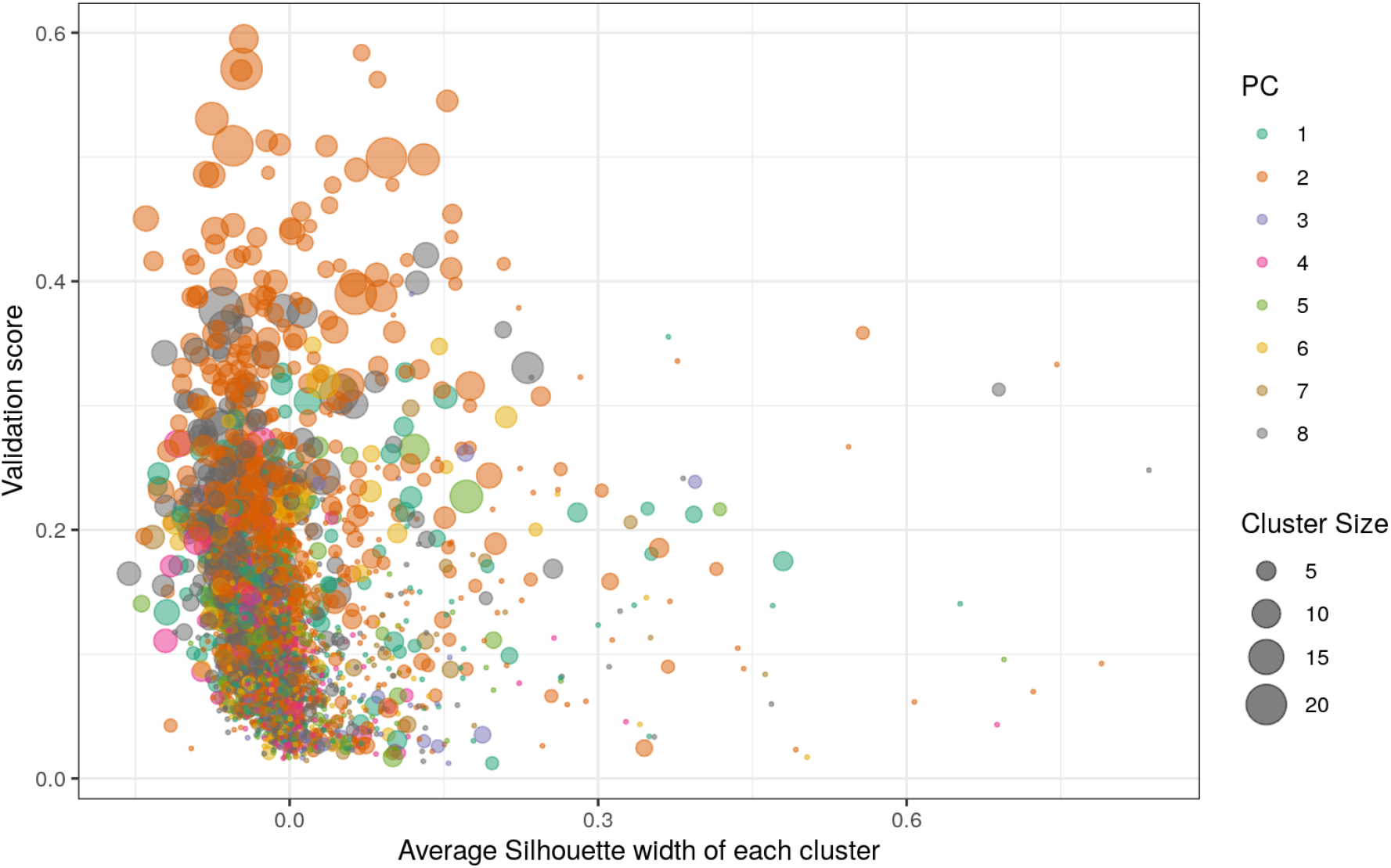
Overview validation results via an interactive plot. In Fig 2B, we used a table format to display the validation results. To understand the overall validation pattern for each PCs of new data, we provide an interactive plot as one of the visualization options. Here, we plotted the validation plot of Human B-cell expression dataset (GSE2350) generated from microarray. X-axis represents the average silhouette width and y-axis represents the validation score. Each point represents RAV, where the color shows the PC with the highest validation score for a given RAV. The point size reflects the cluster size, the number of PCs contributing to a given RAV. In general, we interpret that the points toward the upper right corner with the intermediate sizes are more relevant to new data than the others. An interactive form of this graph is available with the argument `interactive=TRUE`, allowing the user to hover each data point for more information, such as cluster number and exact cluster size.

**Sup.Fig 8.**
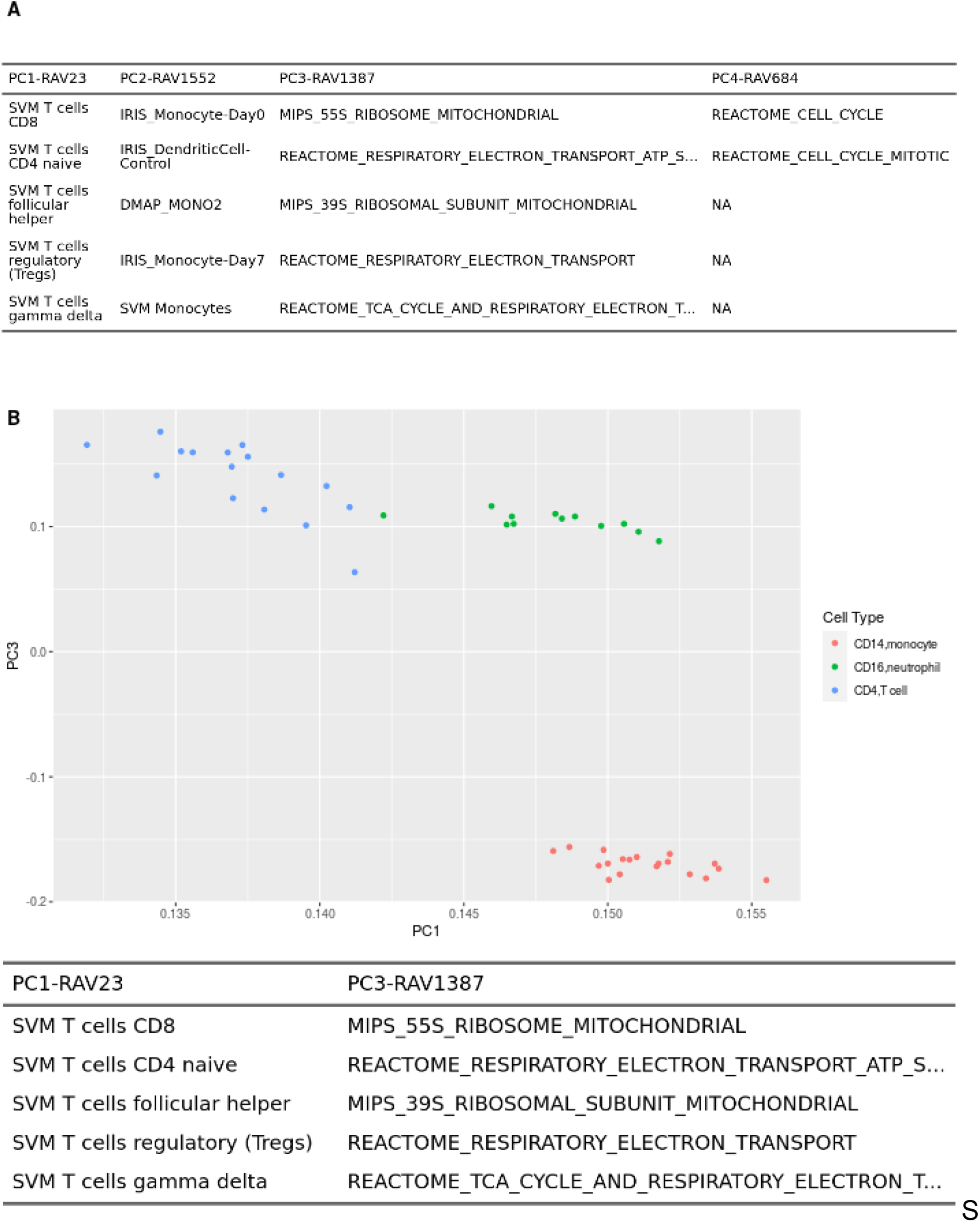
PCA with GSEA annotation. PCA result of leukocyte gene expression data (E-MTAB-2452) is displayed in A) a table or B) a scatter plot. PCA is done on a centered, but not scaled, input dataset by default. Different cutoff parameters for GSEA annotation, such as minimum validation score or NES, can be set.

**Supplementary Figure 9.**
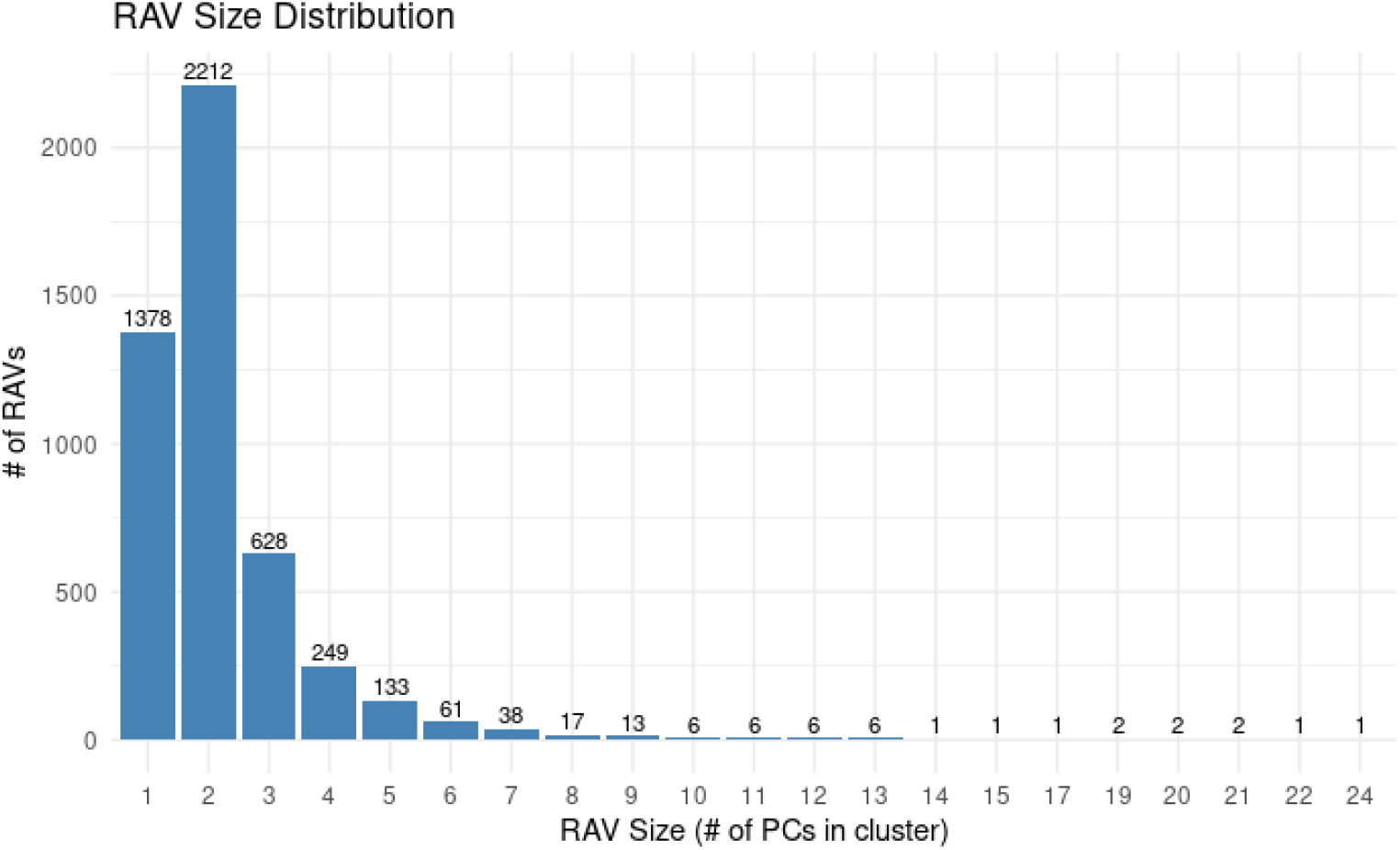
Distribution of RAV sizes. RAVs are constructed from different numbers of PCs, ranging from 1 to 24. Here, we plotted the number RAVs (y-axis) against the cluster sizes (x-axis) to show the distribution of RAV sizes.

**Supplementary Figure 10.**
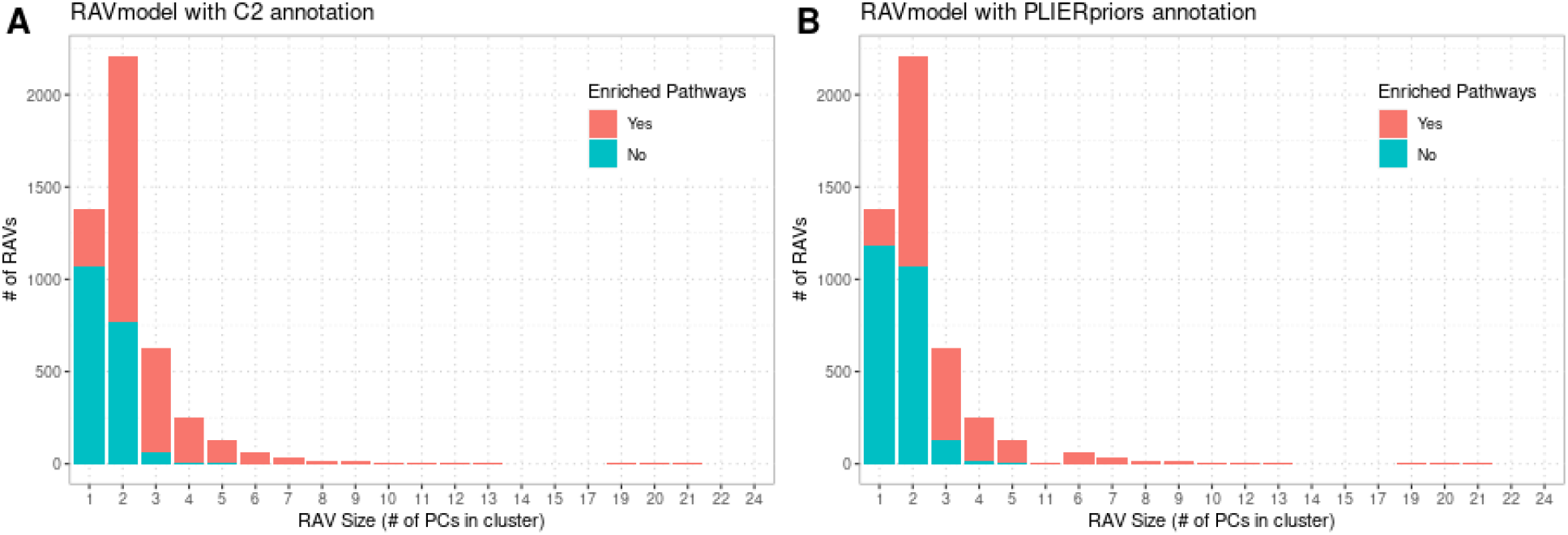
Distribution of PCs in different sized RAVs. https://github.com/shbrief/GenomicSuperSignaturePaper/blob/master/Results/model/PCs_In_Clusters.pdf We summarize the distribution of PCs in different sizes of RAVs. There are 21 different sizes of RAVs with the smallest containing a single PC while the largest containing 24 PCs. We collected all the PCs contributing to the given size of the clusters and plotted 21 barplots. We observed that one- and two- element clusters are predominantly from lower PCs. From three-element clusters, however, the skewness changes from left to right.

**Supplementary Figure 11.** RAVs without enriched pathways. We summarized the gene set annotation status of RAVs based on the RAV sizes. We tested two RAVmodels A) RAVmodel annotated with MSigDB C2 and B) RAVmodel annotated with three gene sets provided through the PLIER package. RAVs without enriched pathways are labeled with teal and RAVs with one or more enriched pathways are in red.

## Supplementary Tables

**Sup.Table 1.**
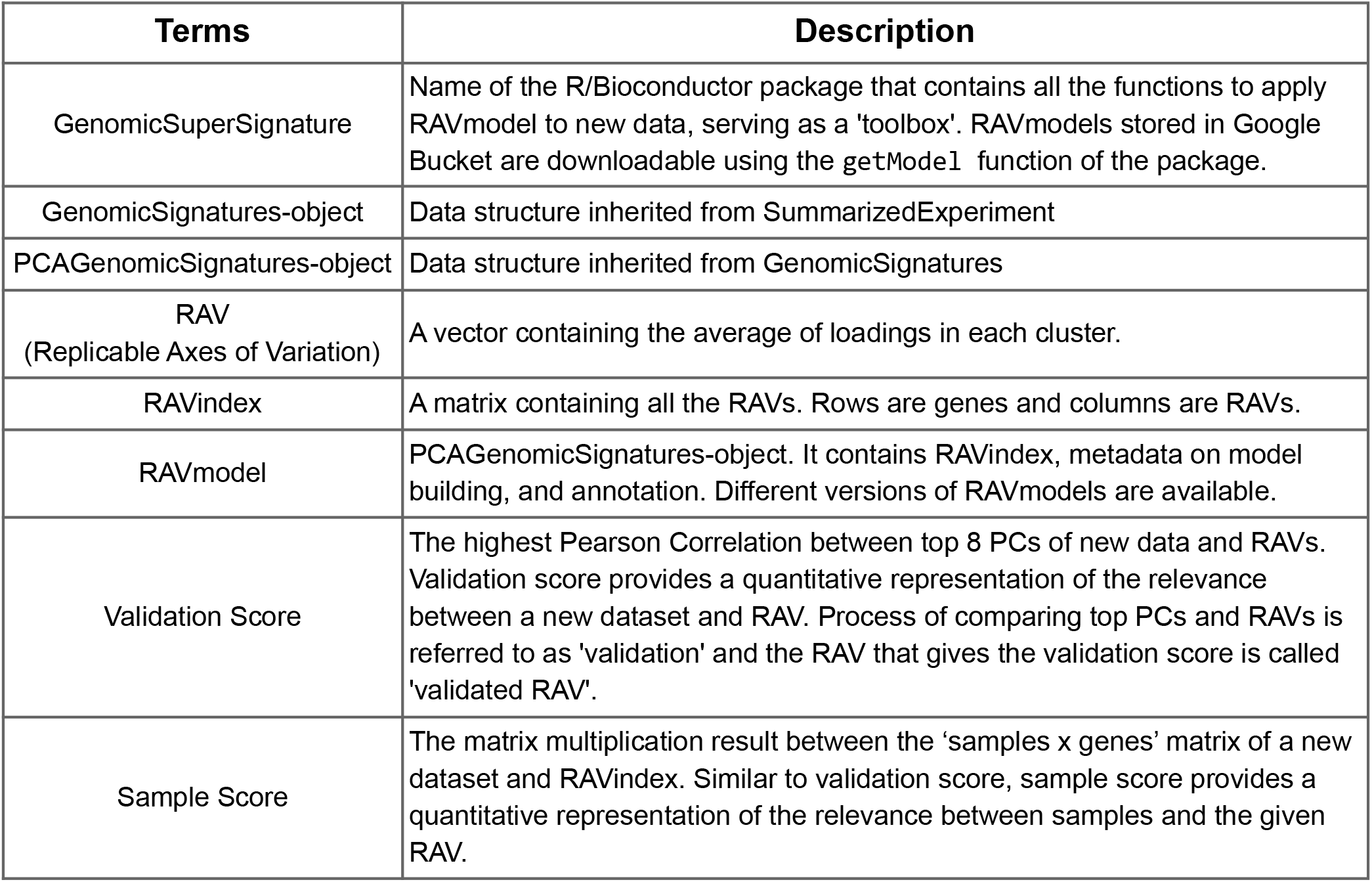
Summary of new terms.

**Sup.Table 2. Training datasets used in this study**

https://github.com/shbrief/GenomicSuperSignature/blob/master/inst/extdata/studyMeta.tsv.gz

The study accession, the number of samples, and the title of 536 training datasets used to construct the current version of RAVmodel (`RAVmodel_536` column). The number of samples for each study based on the metadata (`metadata` column) and the number of samples actually used (`downloaded` column) are different due to the data availability at the time of download.

**Sup.Table 3. Source types of training datasets**

https://github.com/shbrief/GenomicSuperSignaturePaper/blob/master/inst/extdata/source_name_annotated.tsv

We obtained the source name for 435 studies (~81.2% of all training datasets) from OmicIDX^46^ and did a manual curation based on the source name (`source_name` column) to understand the types of training datasets. Curation covered four categories: 1) whether the dataset is cancer or not (`cancer` column), 2) whether the dataset is blood or not (`blood` column), 3) whether the dataset is cell line or not (`cell_line` column), 4) what is the origin of samples (`origin` column).

**Sup.Table 4. Available RAVmodels**

https://github.com/shbrief/GenomicSuperSignaturePaper/blob/master/inst/extdata/SupTable3_RAVmodels.csv

This is the list of currently available RAVmodels that are different in 1) the size of training datasets, 2) the number of top PCs collected from each study, 3) the number of clusters for hierarchical clustering, and 4) gene sets used for GSEA annotation. Two RAVmodels used in this work, RAVmodel_C2 and RAVmodel_PLIERpriors, are available for download using the getModel function and the others are available upon request.

**Sup.Table 5. Summary of top 10 validated RAVs for 18 colorectal cancer datasets**

https://github.com/shbrief/GenomicSuperSignaturePaper/blob/master/Results/CRC/outputs/CRC_top10_validated_ind.tsv

**Sup.Table 6. Summary of boxplot statistics**

https://github.com/shbrief/GenomicSuperSignaturePaper/blob/master/Results/CRC/outputs/boxplot_summary.csv

This is the summary statistics of all the boxplots present in this work. It contains 72 rows from 9 boxplots where each has 4 panels with 2 groups. It has 10 columns labeled as figure, panel, group, minima, whisker_min, first_quartile, median, third_quartile, whister_max, and maxima. Four panel numbers denote MSI status (1), grade (2), stage (3), and tumor location (4). For groups, 1 and 2 represent left and right groups, respectively, in each panel.

**Sup.Table 7.**
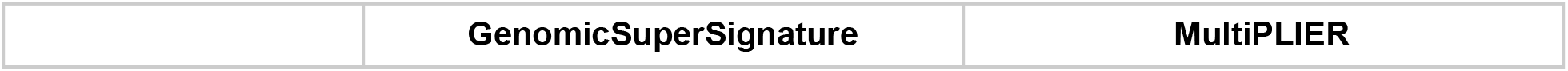
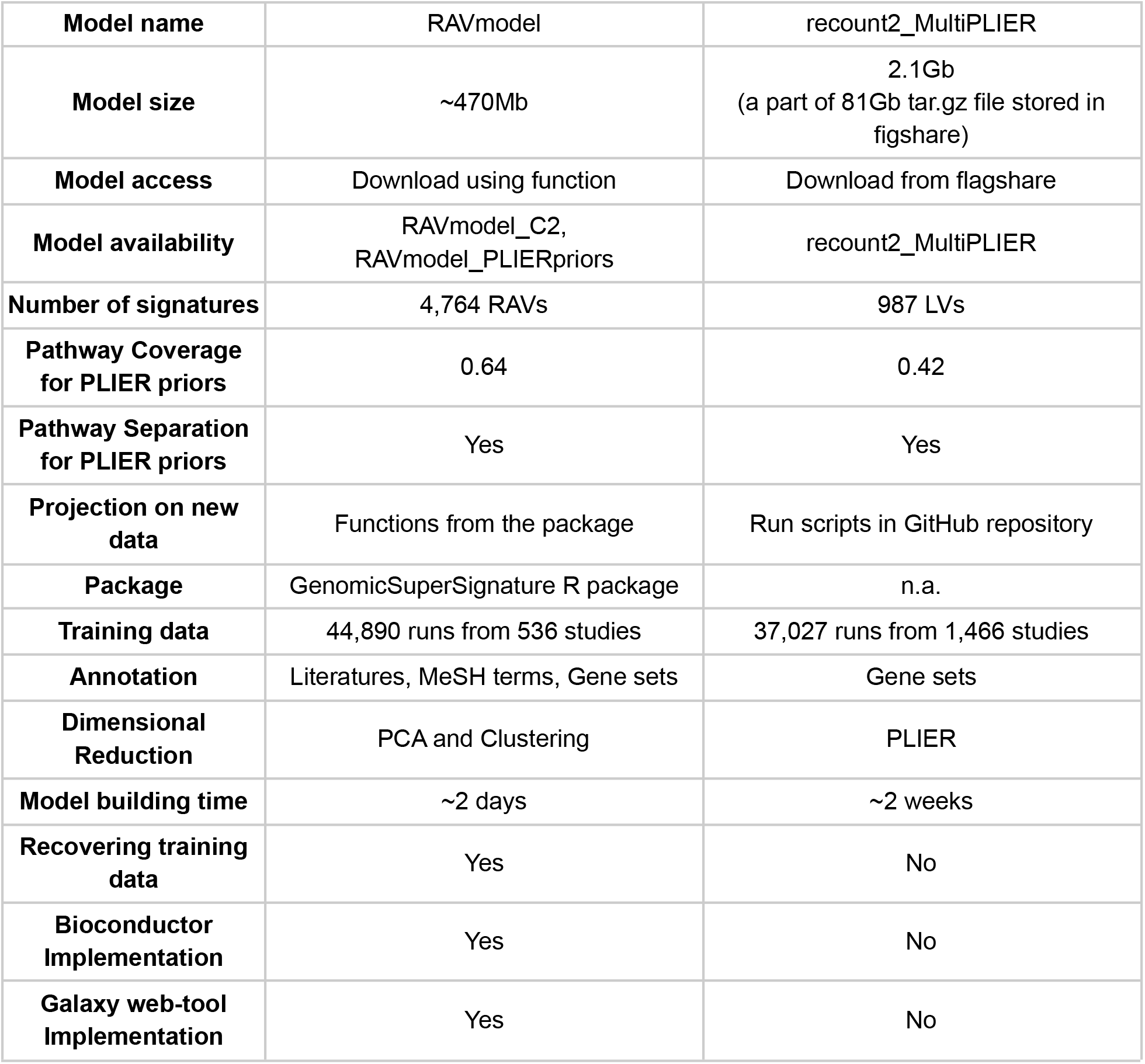
Comparison between GenomicSuperSignature and MultiPLIER.

**Sup.Table 8.**
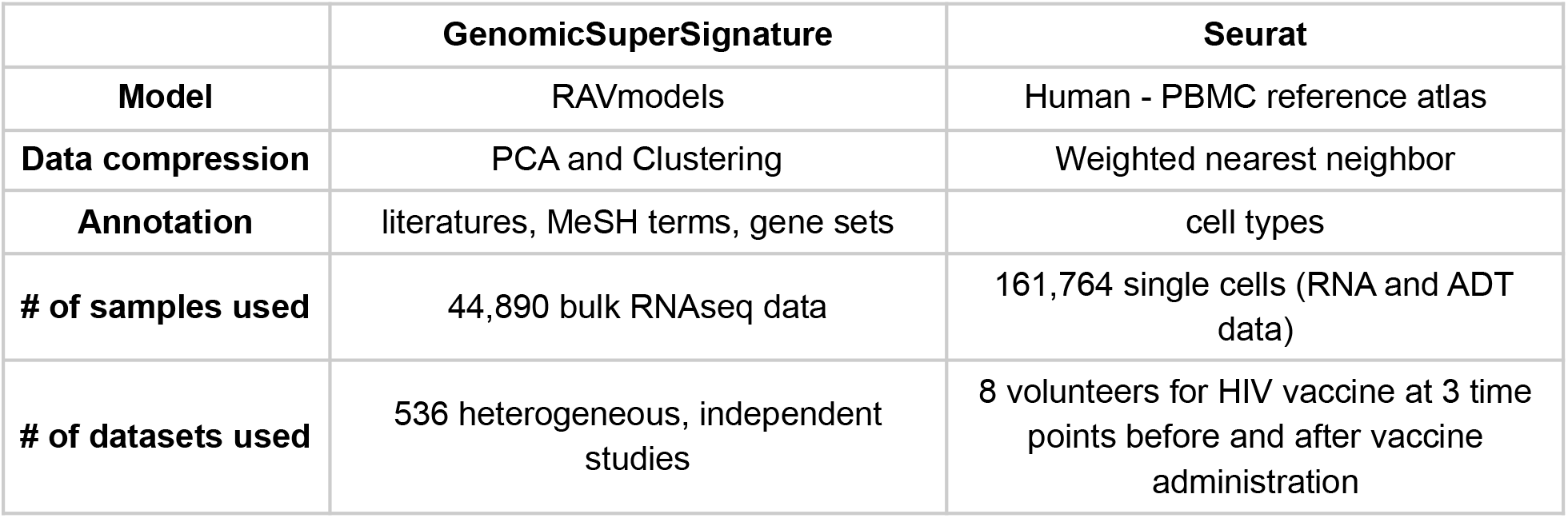
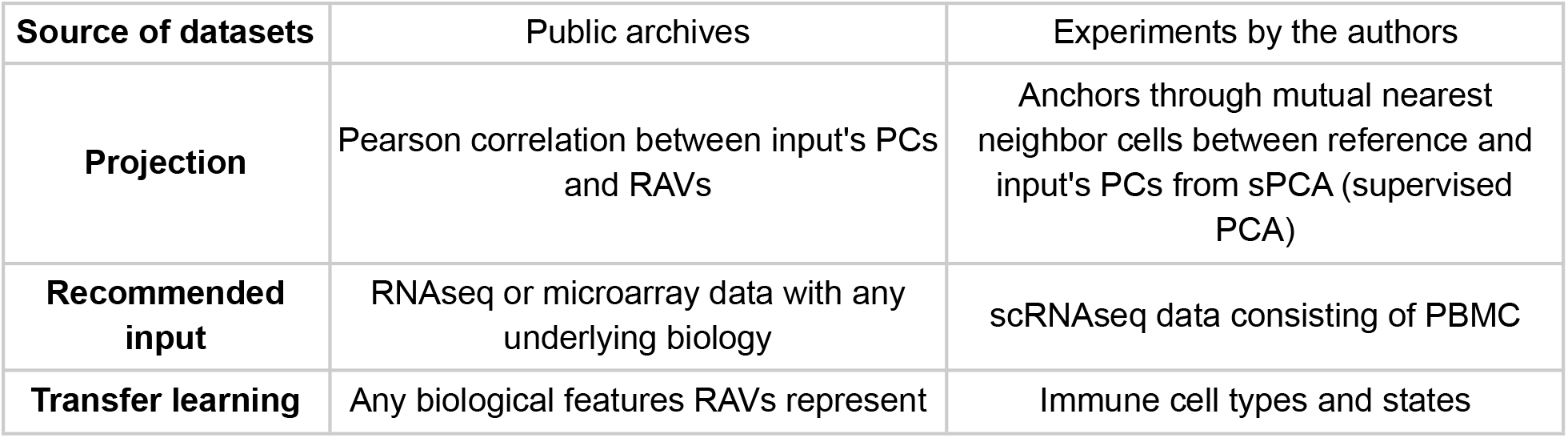
Comparison between GenomicSuperSignature and WNN.

## Notes

http://www.bioconductor.org/packages/GenomicSuperSignature

https://github.com/shbrief/GenomicSuperSignature

### Competing Interest Statement

The authors have declared no competing interest.

